# Common soil history is more important than plant history for arbuscular mycorrhizal community assembly in an experimental grassland diversity gradient

**DOI:** 10.1101/2024.03.14.585138

**Authors:** Cynthia Albracht, Marcel Dominik Solbach, Justus Hennecke, Leonardo Bassi, Geert Roelof van der Ploeg, Nico Eisenhauer, Alexandra Weigelt, François Buscot, Anna Heintz-Buschart

## Abstract

The relationship between biodiversity and ecosystem functioning strengthens with ecosystem age. However, the interplay between the plant diversity - ecosystem functioning relationship and Glomeromycotinian arbuscular mycorrhizal fungi (AMF) community assembly has not yet been scrutinized in this context, despite AMF’s role in plant survival and niche exploration.

We study the development of AMF communities by disentangling soil- and plant-driven effects from year effects. Within a long-term grassland biodiversity experiment, the pre-existing plant communities of varying plant diversity were re-established as split plots with combinations of common plant and soil histories: split plots with neither common plant nor soil history, with only soil but no plant history, and with both common plant and soil history.

We found that bulk soil AMF communities were primarily shaped by common soil history and additional common plant history had little effect. Further, the steepness of AMF diversity and plant diversity relationship did not strengthen over time, but AMF community evenness increased with common history. Specialisation of AMF towards plant species was low throughout giving no indication of AMF communities specialising or diversifying over time. The potential of bulk soil AMF as mediators of variation in plant and microbial biomass over time and hence as drivers of BEF relationships was low.

Our results suggest that soil processes may be key for the build-up of plant community-specific mycorrhizal communities with likely feedback effects on ecosystem productivity, but the plant-available mycorrhizal pool in bulk soil itself does not explain the strengthening of BEF relationships over time.

## Introduction

The relationship between biodiversity and ecosystem functioning (BEF) has been established through manipulation of plant diversity in field experiments (Cardinale *et al*., 2007; Weisser *et al*., 2017; Eisenhauer *et al*., 2019; Hong *et al*., 2022). Especially in temperate grasslands, aboveground plant biomass has been well studied as a proxy for ecosystem functioning (Cardinale, 2011; Eisenhauer, 2012; Tilman, Isbell and Cowles, 2014). The increasing productivity in more diverse communities is commonly observed as overyielding in mixtures in comparison to monocultures (Steudel *et al*., 2016; Weisser *et al*., 2017), of which a large proportion of variation in community biomass is explained by functional diversity (Hector *et al*., 1999; Roscher *et al*., 2012). Plant communities’ adaptations to soil conditions and their interactions with soil biota have been shown to play a central role for productivity (Zuppinger-Dingley *et al*., 2016). In consequence, soil biota have been suggested as main drivers causing the overyielding in more diverse plant communities (Schnitzer *et al*., 2011; Wagg *et al*., 2014; Weisser *et al*., 2017; Eisenhauer *et al*., 2019) and driving belowground facilitation through mutualistic associations (Wagg *et al*., 2015; Wright *et al*., 2017).

Among soil biota, Glomeromycotinian arbuscular mycorrhizal fungi (AMF) are the most promising candidates for drivers of ecosystem functioning. As root endosymbionts, they provide vital nutrients and enable water uptake for ∼80% of all terrestrial plants, they shift competition and adaptation dynamics, enable less competitive species to find their niches and withstand adverse environmental conditions (Smith and Read, 2008; Chowdhury *et al*., 2022; Kuyper and Jansa, 2023). The correlation between plant and AMF diversities has repeatedly been shown (Burrows and Pfleger, 2002; Antoninka, Reich and Johnson, 2011; Guzman *et al*., 2021; Wang *et al*., 2022). Organic matter input by the plant community and increased soil carbon storage over time drive plant-fungal interactions (Lange *et al*., 2014, 2015), with fungi responding to both plant species richness and plant functional diversity (Eisenhauer *et al*., 2017; Lange *et al*., 2023). Vice versa, nutrient uptake through AMF enables different plant species to access different nutrient pools (Wagg *et al*., 2011; Weisser *et al*., 2017). However, the dynamics of a concurrent plant and AM fungal community assembly in the context of BEF relationships remain unresolved.

Prior research has shown that BEF relationships strengthen over time, as a result of complementarity effects (Cardinale *et al*., 2007; Fargione *et al*., 2007; Eisenhauer, 2012; Reich *et al*., 2012; Wagg *et al*., 2022). Complementarity is based on resource partitioning, abiotic and biotic facilitation that lead to selection for specific phenotypes and therefore niche differentiation, which is more pronounced in diverse communities (Tilman and Snell-Rood, 2014; Zuppinger-Dingley *et al*., 2014; Barry *et al*., 2018, 2019). Plant diversity effects on soil microbial biomass likewise develop over time, but with a time-lag as a result of plant-soil feedbacks with accumulation of mutualists in more diverse and antagonists in low diverse communities over time (Habekost *et al*., 2008; Eisenhauer *et al*., 2010; Eisenhauer, 2012; Thakur *et al*., 2015; Strecker *et al*., 2016). This plant-soil feedback is further driven by resource availability like soil carbon (C) and soil nitrogen (N) storage, which is increased at increased root biomass and microbial activity in more diverse communities (Fornara and Tilman, 2008; Lange *et al*., 2015).

As AM symbiosis is a result of its ecological context shaped by biotic factors like plant root architecture and efficiency of fungal nutrient turnover, as well as abiotic factors like soil properties, all of which are dynamic, we would expect that AM symbiosis and consequently AM fungal communities are also a product of temporal dynamics (Ji and Bever, 2016; Rudgers *et al*., 2020; Berger and Gutjahr, 2021). In fact, the long-term biodiversity research platform of the Jena Experiment, started in 2002, has provided a moving image of these relationships. In the Jena Experiment, plant species richness (1-60 native species) and functional richness (4 functional groups) are as independently as possible manipulated in the 80 maintained grassland plots. AMF communities analysed in 2007 were significantly affected by both plant species richness and plant functional richness (König *et al*., 2010a). However, in 2010, plant species richness only marginally positively affected total fungal community diversity and had no significant impact on fungal community composition (Dassen *et al*., 2017). In 2017, a weak positive relationship between plant and total fungal, but not AMF diversity was observed (Albracht *et al*., 2023), while plant functional group composition significantly affected AMF and total fungal community composition (Albracht *et al*., 2023; Maciá-Vicente *et al*., 2023).

As shown by this plant and fungal diversity relationship alone, effects change with calendar year and it is difficult to separate actual age effects from short-term variations caused by e.g. weather conditions or general trends of climate change as well as community assembly. Climate events such as droughts or floods have been shown to influence both plant productivity and BEF relationships (Vogel, Scherer-Lorenzen and Weigelt, 2012; Wright *et al*., 2015; García-Valdés, Bugmann and Morin, 2018). The ΔBEF experiment (**Fig. 1**) was nested within the Jena Experiment in 2016 to disentangle year effects from history effects by re-establishment of the original biodiversity experiment 14 years after the initial establishment of the experimental communities (Vogel *et al*., 2019). ΔBEF contains three treatments of 1) re-established split plots with no common plant and soil history (NH), 2) split plots with new plants of the original composition on maintained soil with community-specific history (SH), and 3) the maintained plots with common plant and soil history (PSH). By maintaining the common plant and soil histories of these 14-year-old grassland communities or re-establishing plants or plants and soil, the ΔBEF experiment not only separates BEF relationships from the respective calendar years with their conditions, but also allows to test the individual part of plant and soil history in the age effect. Note that we use the term ‘common history’ rather than ‘legacy’ throughout, because ‘common history’ makes no assumption on whether the newly applied plant community signifies a regime shift while the term ‘legacy’ indicates that we observe remnants of previous plant communities despite the effects of a new community.

**Fig. 1.**
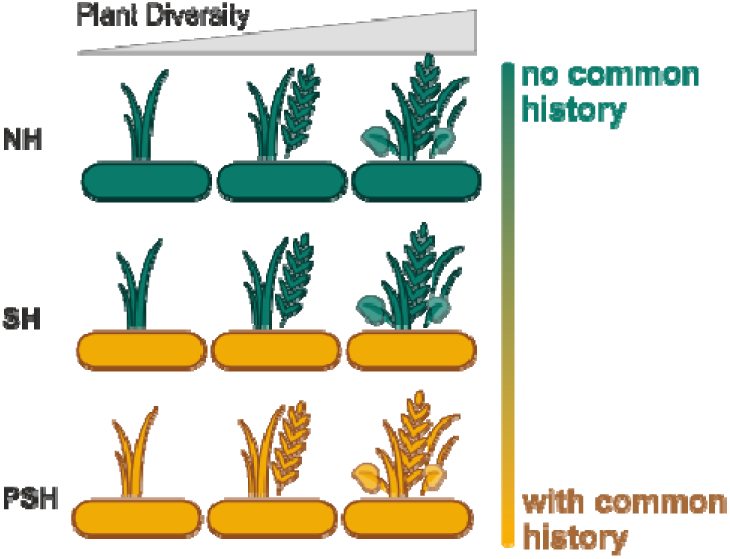
Simplified design of the ΔBEF experiment sampled here: Plant communities with sown diversity ranging from monocultures, over 2, 4, 8, to 16 species mixtures, varying in composition and functional diversity (14-16 communities per diversity level). In split plots with no common history (NH) plant material and topsoil were removed and split plots were re-sown 5 years prior to sampling; SH split plots kept original soil and only had the plant communities re-sown 5 years before sampling; the split plots with common plant community and soil history (PSH) remained from the main Jena Experiment established 19 years prior to sampling.

In this setup, Macía-Vicente et al (2023) could show that while the relationship of fungal diversity and plant diversity was weaker in split plots sharing 15 years compared to 1 year of common plant history, the fungal community composition showed an increasing association with plant diversity over time. The same analysis found that the relative abundance of AMF among the fungal community decreased over time, but AMF diversity was more strongly related to plant diversity after 15 years compared to 1 year of common soil history, suggesting a gradual shift towards specific AMF assemblages (Maciá-Vicente *et al*., 2023). However, because of the use of universal fungal primers, the resolution of AMF in this study was limited. In addition, the comparison was restricted to split plots with the original plant community and re-established plant communities, and no experimental re-establishment of the soil community was analysed.

With the present study, we investigated how species-rich and -poor plant communities with common plant and soil histories affect AMF communities and vice versa. Thereby, we explored whether plant or soil histories are more important drivers of AMF community composition and diversity. Continuing from the work of Macía-Vicente et al (2023), we sampled bulk soil at a later stage of the ΔBEF experiment with focus on AMF using specific primers. We further expanded the analysis to all combinations of grassland communities of 19 vs 5 years of common plant and soil history, therefore also included the soil history as a factor of developing BEF relationships.

Given the steeper BEF relationship in plant communities with established soil history, we expected (H1a) plant communities with long-term common soil history to have stronger effects on AMF community composition (comparison SH vs NH). In this context, AMF community composition should respond to plant community diversity and composition. Given the tight interaction between AMF and plants, and in contrast to previous results on other fungal guilds (Maciá-Vicente *et al*., 2023), we further expected (H1b) common plant history to lead to additional strong effects on AMF community composition (comparison PSH vs SH). We anticipated AMF diversity to increase with increasing plant diversity in accordance with the abovementioned previous results and hypothesised that (H2) this relationship strengthens with common soil history and – to a lesser extent – additional common plant history. Complementary to (H2), we expected (H3a) AMF to shift from mainly generalists to more specialists in grasslands with common history, therefore diversifying and exploring more niches after the grassland communities have been established. At the AMF community level, this would lead to (H3b) more divergence in contrasting plant communities in split plots with common history. And finally, with AMF being potential drivers of BEF relationships, we expected (H4) AMF communities to drive plant productivity more in grasslands with common history.

## Material and Methods

### Field Experimental Design

As the basic setting, we used the existing long-time biodiversity experimental platform of The Jena Experiment in Jena, Germany, which consists of 80 grassland plots of differing plant diversity (1, 2, 4, 8, 16, or 60 grassland species) and functional composition (4 functional groups: small and tall herbs, legumes, and grasses) (Roscher *et al*., 2004). These 80 plots of varying composition out of a 60 grassland species pool have been maintained since 2002.

In 2016, the ΔBEF experiment was established with a setup of three history treatments with varying age of the plant and soil communities, as described in detail in Vogel et al (2019) (**Fig. 1**). In the first treatment, arable soil from an adjacent crop field was filled into excavated 1.5 x 3 m split plots, and the plot-specific plant communities were sown from unrelated seeds (Vogel *et al*., 2019). This treatment is hereafter called NH (<u>n</u>o <u>h</u>istory) as plants and soil have no common history. The soil does however resemble the Jena Experiment soil at the beginning of the original experiment (Vogel *et al*., 2019). For the second treatment, the soils from the pre-existing experiment plots were kept, but plant material was removed from split plots, and new plants were sown (again from unrelated seeds). The split plots of this second treatment, hereafter called SH (only soil <u>h</u>istory) therefore contained soils that has had a long time to adapt to its specific plant community, but the newly sown plants had no history with the soil. The third treatment was formed by the pre-existing plots of the Jena Experiment with old soil and plant communities (PSH, with <u>p</u>lant community history and with soil <u>h</u>istory).

Soil of all plots with monocultures or 2- to 16-species mixtures (i.e., 14-16 plots per diversity level) and all treatments (i.e., 3 split plots per plot) was harvested between May 31^st^ and June 12^th^ 2021. At this point, the newly established plant communities and soil had existed for 5 years, and the original soil and communities for 19 years. The 5 years passed should have allowed the NH split plots to recover from the disturbance (Schmid *et al*., 2021). For AMF community analysis on plot level, 4 soil cores (5 cm depth, 4 cm in diameter) were taken per split plot. One combined sample of bulk soil per split plot was frozen at -20°C until further processing.

### AMF marker gene sequencing and data processing

Genomic DNA was extracted from bulk soil samples using Quick-DNA™ Fecal/Soil Microbe Miniprep Kit (Zymo Research). Quantity and purity of DNA isolated from bulk soil samples were assessed using a NanoDrop 8000 spectrophotometer (Thermo Fisher Scientific). The DNA was then amplified in triplicates. We targeted the SSU rRNA region of AMF with primer pairs Glomer1536 and WT0 (AATARTTGCAATGCTCTATCCCA / CGAGDWTCATTCAAATTTCTGCCC (Wubet *et al*., 2006; Morgan and Egerton-Warburton, 2017) for the first and NS31 and AML2 (TTGGAGGGCAAGTCTGGTGCC / GAACCCAAACACTTTGGTTTCC (Simon, Lalonde and Bruns, 1992; Lee, Lee and Young, 2008) for the second PCR step following the protocol of Wahdan et al (2021). The samples were prepared for Illumina sequencing by purifying amplicons, followed by barcoding and quality checking per the protocol of Wahdan et al (2021). The sequencing was performed at the Illumina MiSeq platform at the Department of Soil Ecology of the Helmholtz Centre for Environmental Research – UFZ, Halle (Saale), Germany as paired-end sequencing of 2 x 300 bp.

The sequencing output was processed with the *snakemake* implementation of *DADA2* (Callahan *et al*., 2016) *dadasnake* (Weißbecker, Schnabel and Heintz-Buschart, 2020): primers were trimmed and sequences filtered to a minimum length of 260 bp (fwd) / 210 bp (rvs); reads with expected error higher than 2 were discarded; amplicon sequencing variants (ASVs) were determined with ‘pooled’ settings and forward and reverse ASVs were merged with a minimum overlap of 12 bp and 0 mismatches and filtered for chimaeras. Taxonomic classification was done with *mothur* (Schloss *et al*., 2009) against the SILVA v138 SSUref database (Quast *et al*., 2013) to then discard non-Glomeromycotinian ASVs.

The remaining fungal ASV were blasted against the MaarjAM database (Öpik *et al*., 2010) to be assigned to virtual taxa (VTX). All ASVs that could not be assigned to a virtual taxon were extracted, filtered for singletons and used to construct a maximum likelihood phylogenetic tree based on a general time-reversible, discrete gamma (GTR+G) model using raxML (Stamatakis, 2014) and FasttreeMP (Price, Dehal and Arkin, 2010). Thus, these ASVs were assigned custom virtual taxa (VTC) with cophenetic distances below 0.03 (Albracht *et al*., 2023).

### Soil and plant data

Aboveground plant biomass was harvested from May 31^st^ to June 8^th^ 2021 and measured in two 0.1 m^2^ (20 x 50 cm) frames per split plot as described in Vogel et al (2019). Briefly, plants were cut with scissors at around 3 cm above soil surface and stored in plastic bags at 4°C until they were sorted into target (sown) species, weeds (not sown), rest (unidentifiable plant material), and litter (dead plant material). The sorted samples were weighed after drying at 70°C for at least 48 h. Final biomass is given in g/m^2^ based on the sampled area minus bare ground area.

From May 31^st^ to June 11^th^ 2021, an additional 4 soil cores of 5 cm depth and 3.5 cm diameter were taken in a strip of 1 m in the centre of each split plot. These were bulked together per split plot and all roots washed and separated into coarse (> 2 mm) and fine (< 2 mm) roots to determine root biomass. All roots were dried at 70 °C for at least 48 h and then weighed. The final root biomass is given in g/m^2^ based on the calculated sampled area: *number of cores * (1.75 cm)^2^ * π*.

For soil phosphorus (P), about ¼ of the soil cores used for root biomass determination was combined, sieved to < 2 mm and dry-weight equivalents of 1 g sample material were extracted with a 0.5 M NaHCO_3_ solution (Carl Roth) at pH 8.5 for 30 min following Olsen (1954). PO_4_ concentrations were analysed after filtering extraction solution (Mn 619 G1/4, Macherey-Nagel) using continuous flow analyser (Seal Analytical) with molybdenum-blue method (Murphy and Riley, 1962). Soil water content (SWC) was analysed gravimetrically from the same soil samples with a halogen moisture analyzer (HB43-S Halogen, Mettler Toledo).

Samples for soil microbial respiration were taken at the end of June 2021, pooling 4 soil cores per split plot of 10 cm depth and a diameter of 2 cm which were stored at 4°C until analysis. The soil was sieved (<2 mm) and visible plant material and animals removed. The analyses were performed with an O_2_ micro-compensation apparatus measuring basal respiration (BR), maximum initial respiratory response (MRR), soil microbial carbon biomass (Cmic) and specific respiratory quotient (qO2).

### Statistical analysis

All statistical analyses were done in *R version 4.1.1* (R Core Team, 2017), unless stated otherwise. Virtual taxa (VT, including VTX and VTC) were filtered to at least one count in a minimum 2 % of all samples.

We performed an explorative PARAFAC analysis (Carroll and Chang, 1970; Harshman, 1970; Bro, 1997) on filtered and rarefied abundance data which was exported to *MATLAB vers. 9.14.0* (The MathWorks Inc., 2022). In MATLAB, the abundance data was cubed, a pseudocount of 1 added and the data centre-log transformed. To perform a PARAFAC analysis (Bro, 1997) based on the *N-way toolbox* (Bro, 2023), the data was centred across VT and scaled within VT (Bro and Smilde, 2003), before using bootstrapped initiation of 100 PARAFAC models and selecting the number of valid components based on steady CORCONDIA values (100) across the initiations. The PARAFAC outputs were exported back to R for summarization and averages of bootstrapped models plotted with *ggplot2* (Wickham, 2009).

To explore differences in β-diversity across the history treatments, Aitchison distances were calculated with package *robCompositions* (Templ, Hron and Filzmoser, 2011) from the filtered and rarefied abundance matrix after replacing zeros with a pseudo-count of 0.1. All pairwise distances were extracted and compiled for each treatment and plant diversity combination. We used Wilcoxon signed rank tests on the pairwise distances to find significant differences in beta-diversities across the history and diversity gradient.

To quantify history treatment and plant diversity effects on AMF communities, we ran PERMANOVA (*adonis2* function in *vegan*, (Oksanen *et al*., 2019)) on the Aitchison distance matrix (*distances ∼ history treatment * plant diversity _log_* (9999 permutations, stratified for block)). As we and others have previously observed plant functional group presence to play an additional role in shaping AMF communities (König *et al*., 2010b; Albracht *et al*., 2023; Maciá-Vicente *et al*., 2023), we further ran a more complex PERMANOVA on the Aitchison distance matrix to test for effects of plant community compositions:

*distances ∼ history treatment + plant diversity _log_ + functional diversity + legumes _p/a_ + grasses_p/a_ + herbs_p/a_*.

Barplots were based on relative abundances calculated in *phyloseq* (McMurdie and Holmes, 2013) on the filtered abundance data. We used *Maaslin2* (Mallick *et al*., 2021) with a zero-inflated negative binomial (ZINB) model on the count data normalised to trimmed mean of M-values (TMM) to calculate differential abundances. We ran three individual models to find differentially abundant VT between history treatments (plot as random effect), between the plant diversity levels (block and treatment as random effects), and the combined model of history treatment and plant diversity and their interaction (block and plot as random effect). For the second model, we tested both a regression model with VT read counts on the logarithm of plant diversity and, to detect nonlinear plant diversity effects, a categorical model comparing higher diversity levels against the monocultures.

□-diversity indices (Richness, Shannon and Simpson diversity indices, Pielou’s evenness) were estimated with *phyloseq* (McMurdie and Holmes, 2013) on rarefied data (2400 reads per sample). The □-diversity indices were tested with the linear mixed model:

*□-diversity ∼ history treatment * plant diversity_log_.*

As the Jena Experiment field site has a known edaphic variation with *soil phosphorus* (P) and *soil water content* (SWC) varying across the 4 blocks the grassland plots are grouped in, the block and plot were carried as a random effect.

We assessed whether AMF VT increase or lose plant partner specificity from younger to older grasslands. The degree of specialisation of AMF VT to plant species was calculated as the phi-coefficient (Chytrý *et al*., 2002; Weißbecker *et al*., 2019) per treatment:

*φ = ± √(Χ ^2^/N) = (a × d-b × c)/√((a + b) × (c + d) × (a + c) × (b + d))*

where *a* is the number of occurrences of a VT in a split plot where a particular plant species was sown, *b* the number of occurrences of the VT in split plots without that plant, *c* the number of times the VT is absent in split plots where the plant was sown, and *d* the number of times the VT is absent in split plots without the plant. The resulting *φ*-coefficients range from -1 to 1 with values above 0 indicating a specificity between the individual VT and plant species. Significance was determined by comparison to null-models. We used 1000 simulations of a *c0* model (preserving VT frequencies) and a *greedyqswap* model (additionally preserving VT richness per sample) in *vegan* (Oksanen *et al*., 2019) to generate FDR-adjusted p-values for the significance of the *φ*-coefficients.

We analysed species turnover on filtered and rarefied count data with package *codyn* (Hallett *et al*., 2016) calculating appearances, disappearances, and the total turnover from treatments NH to SH and SH to PSH.

The edaphic variables were tested with ANOVA against the history treatment and the plant diversity (*env ∼ history treatment * plant diversity _log_*) and in a second model against the diversity and presence/absence of functional groups to test how composition and age of the plant communities interact with soil parameters and plant productivity. The edaphic data further was used for mediation analysis with *LDM vers. 5* (Hu and Satten, 2022) to analyse which changes in environmental factors are potentially mediated by the AMF communities (base model: VT_abundance_table | confounder ∼ (factors) + (target e.g. aboveground biomass) with confounder = 1 and three different models in which the factors were a. (history treatment + plant diversity_log_ + soil P + SWC) or only b. (history treatment + plant diversity_log_) or c. (soil P + SWC).

## Results

### Exploration of the AMF community in plant diversity and history treatments

We obtained a total of 3,967,994 reads (2,450 to 32,249 reads per sample) assigned to *Glomeromycota* ASVs. The ASVs were aggregated to 128 VT, of which 106 were defined by alignment to the MaarjAM database (VTX; Online Resource 1: **Table S1**) and 22 from a de-novo maximum likelihood phylogenetic tree (VTC; Online Resource 1: **Table S2**). The majority of AMF in the bulk soil belonged to the genera *Glomus* (46.6 %) or *Claroideoglomus (* 24.5 %), followed by *Diversispora* (overall 18.5 %). The custom clusters (VTC) had lower relative abundances ranging from < 1 to 4.3 % (Online Resource 1: **Fig. S1**).

We explored patterns of rarefied AMF abundances across the plant diversity and history treatment using PARAFAC three-way analysis. The best model described a single component with an explained variance of 6.56 % (+-0.15 %), demonstrating a weakly reliable relationship between AMF, plant diversity and history treatment. VT abundances responded to the plant diversity gradient with the relationship increasing with plant diversity (**Fig. 2a**). VT determined as *Claroideoglomus* responded more to high-diverse plots (**Fig. 2b**), while VT of genus *Glomus* – while showing diverse patterns – displayed the opposite relationship. Out of the history treatments, PSH displayed a slightly more pronounced VT-plant diversity relationship (**Fig. 2c**).

**Fig. 2.**
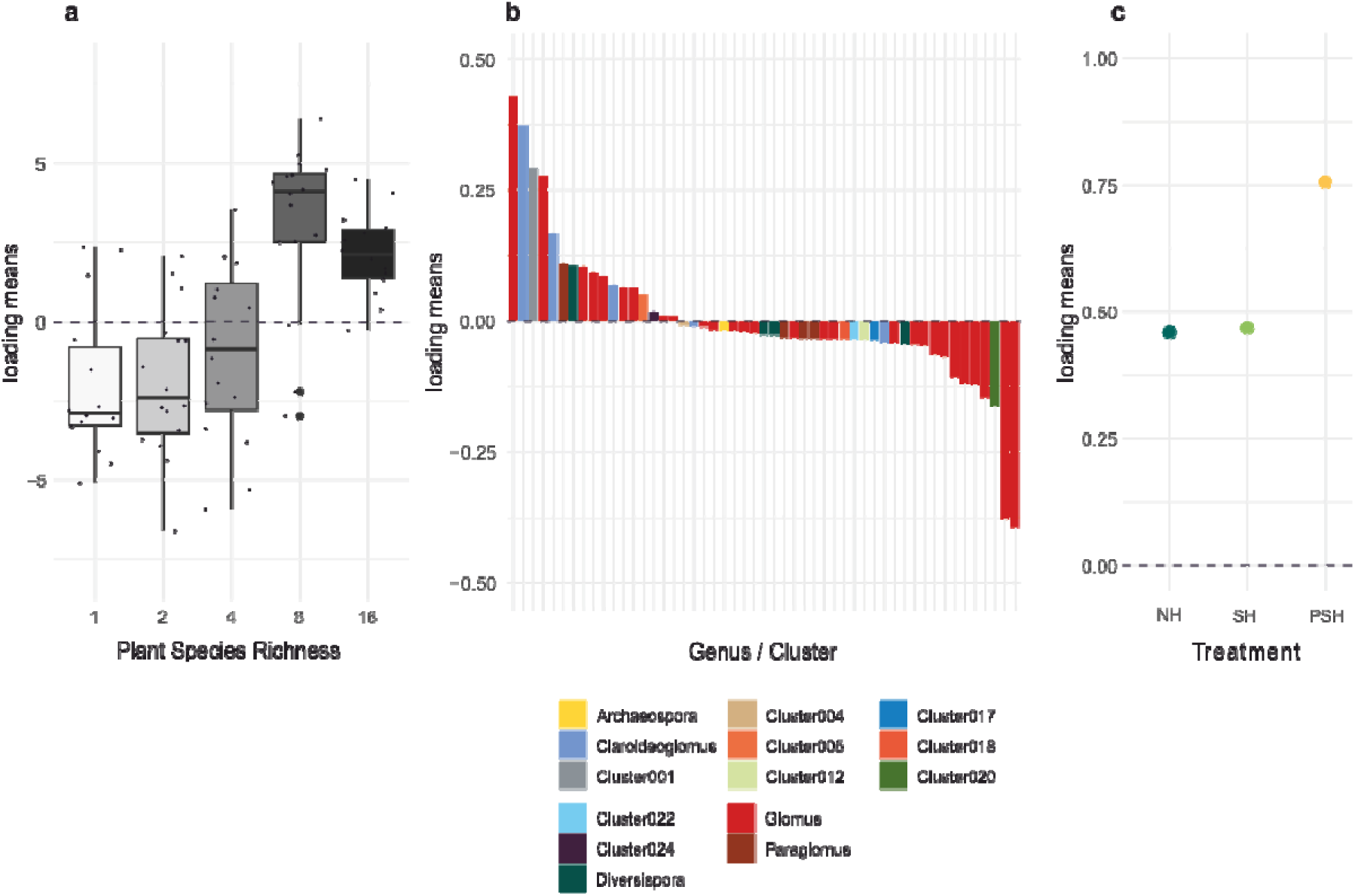
Three-way exploration of VT abundances across plots and history treatments. **(a)** Plot mode: across the data cube, the plant diversity gradient had a strong influence on VT abundance. **(b)** VT mode: in the context of the modes displayed in the other panels, loadings >0 of VT indicate a higher response of this VT in more diverse plant communities and lower response in low-diverse plant communities; loadings < 0 of VT indicate the opposite trend. **(c)** Treatment mode: the history treatments all follow similar trends (all loadings > 0) with the VT and plant diversity interaction being most pronounced in PSH (with plant community and soil history). Means of loadings from 100 bootstrapped PARAFAC analyses explaining 6.56% of the variance are displayed.

### Plant diversity and identity effects in split plots with different histories

In accordance with the exploratory analysis, a PERMANOVA against the full Aitchison distance matrix revealed that history treatment had a significant, but low impact, on AMF communities (R^2 = 0.09, p < 0.001). Testing pairs of history treatments with PERMANOVA, we found that the history treatment effect was larger for soil history (history treatment effects NH vs SH and NH vs PSH: R^2 = 0.09, p < 0.01) than for the plant history (SH vs PSH: R^2 = 0.02, p < 0.01).

Accordingly, when taking all plant diversity levels together, the AMF communities in PSH split plots were significantly more distant from NH communities than from SH communities (**Fig. 3a**). NH split plots were as different from PSH as SH split plots. Hence, soil history, which differentiates SH and PSH from NH split plots, seems to be the more important driver shaping AMF communities in this grassland. However, the greater similarity of PSH and SH split plots was only observed in plant communities with 4 or more species (**Fig. 3b-f**).

**Fig. 3.**
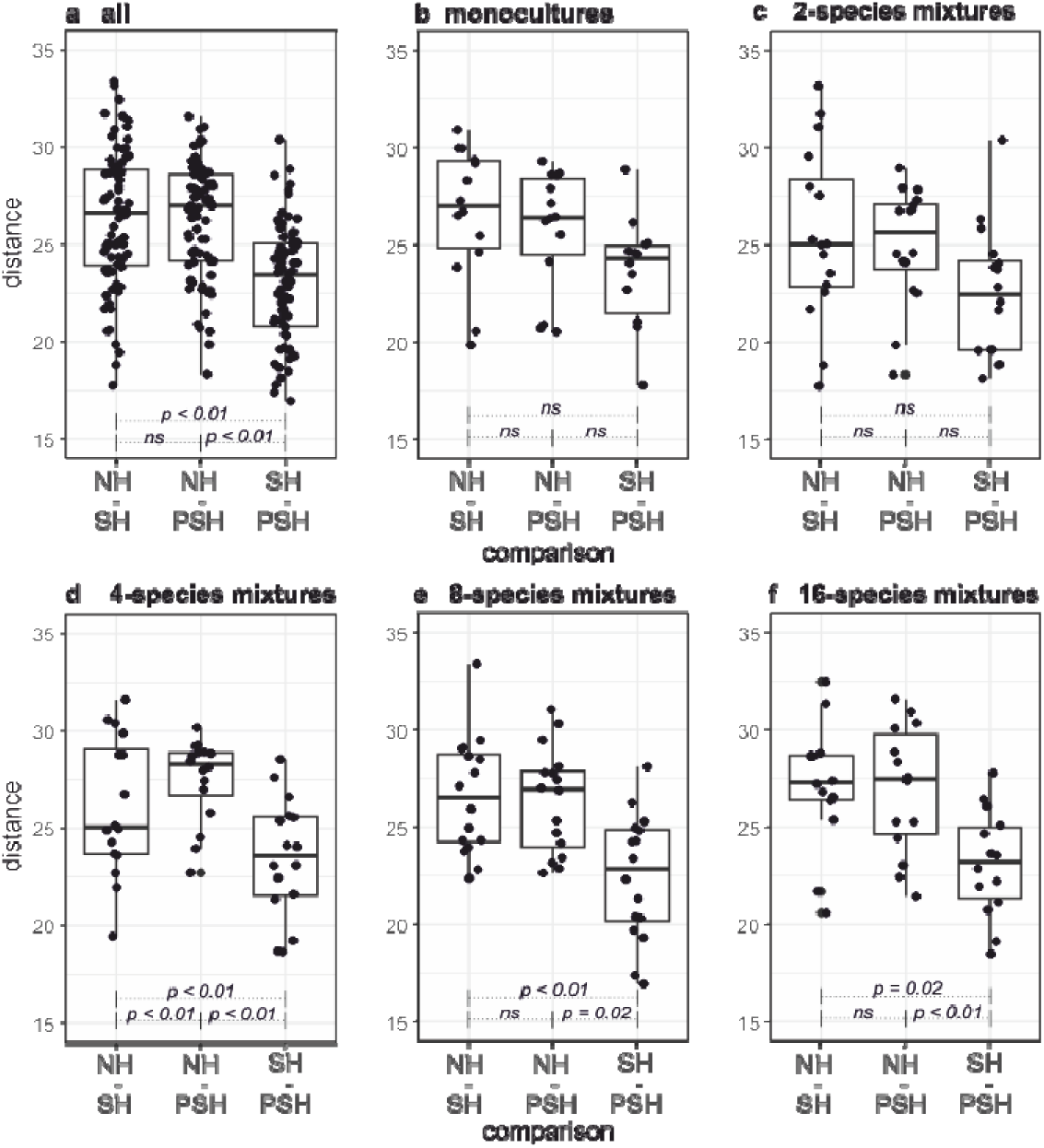
Comparisons of pairwise Aitchison distances of AMF communities between history treatments in the **(a)** monocultures, **(b)** 2-species plots, **(c)** 4-species plots, **(d)** 8-species plots, and **(e)** 16-species plots. P-values indicate significant differences of the distances between history treatments (Wilcoxon’ signed rank test), n.s: not significant at alpha=0.01.

Plant diversity (R^2 = 0.02, p < 0.001) and the interaction of history treatment and plant diversity shaped AMF communities (R^2 = 0.01, p < 0.01), however both with lower effect sizes. The plant diversity effect was not stronger in NH (R^2 = 0.03, p = < 0.01) than SH (R^2 = 0.02, p < 0.01). The plant diversity effect was however stronger in PSH split plots (R^2 = 0.06, p < 0.001) than in either split pot without plant community history, indicating that plant diversity effects on AMF communities depend on shared plant community history.

We tested additional potential drivers of AMF community composition among plant community composition. We ran a PERMANOVA on the AMF community composition against the history treatments and plant diversity by adding the number of functional groups and presence/absence of individual plant functional groups. This revealed that the number of different functional groups (R^2 = 0.01, p = 0.37) was not important, however presence of functional plant groups explained some variation (legumes: R^2 = 0.01, p < 0.001; grasses: R^2 = 0.01, p = 0.04; herbs: R^2 = 0.01, p = 0.01). Relating these findings to edaphic factors, soil P was significantly lower when legumes (t = - 2.99, p < 0.01) were present and significantly higher with grasses (t = 2.23, p = 0.03) present. Zooming in on edaphic differences, NH split plots contained significantly higher P (F = 65.22, p < 0.01) and had a significantly higher SWC (F = 4.19, p = 0.02) than the split plots with soil history. SWC was higher with grasses present (t = 2.12, p = 0.04). The presence/absence of herbs had no influence on either soil P (t = -0.84, p = 0.40) or SWC (t = 0.00, p = 0.99).

### Differences in relative abundances of VT across history and plant diversity

As we found AMF communities to shift with soil history and plant diversity, we calculated differential abundances to find which taxa contributed most to these shifts and to discern whether history treatments induced similar or differing changes. Most VT showed the same direction of change in SH and PSH compared to NH (**Fig. 4a**) and there was no clear trend in PSH having an additional effect to SH. The strongest difference according to soil history was found for VTX00193 (*Claroideoglomus*), which was significantly more abundant in split plots with soil history (**Fig. 4a**). The same trend could be seen for another *Claroideoglomus* VTX00340. Other *Claroideoglomus* (VTX00056, VTX00357) were only higher in SH split plots, but not in PSH. Among the *Paraglomus,* virtual taxon VTX000281 was less abundant in both SH and PSH than in NH, while VTX00444 and VTX00335 were less abundant in only SH or PSH compared to NH. The genus *Diversispora* had mixed results with VTX00054 being less abundant in SH, while VTX00380 was higher in PSH and VTX00062 higher abundant in both SH and PSH in comparison to the NH split plots. *Cluster 20*, which likely contains *Diversispora,* was more abundant in split plots with soil history, as well. *Glomus* also showed diverse responses to the history treatment with e.g. VTX00214, VTX00130 and VTX00153 being more abundant in split plots with soil history, whereas VTX00155, VTX00166 and VTX00064 were significantly less abundant in the split plots with soil history.

**Fig. 4.**
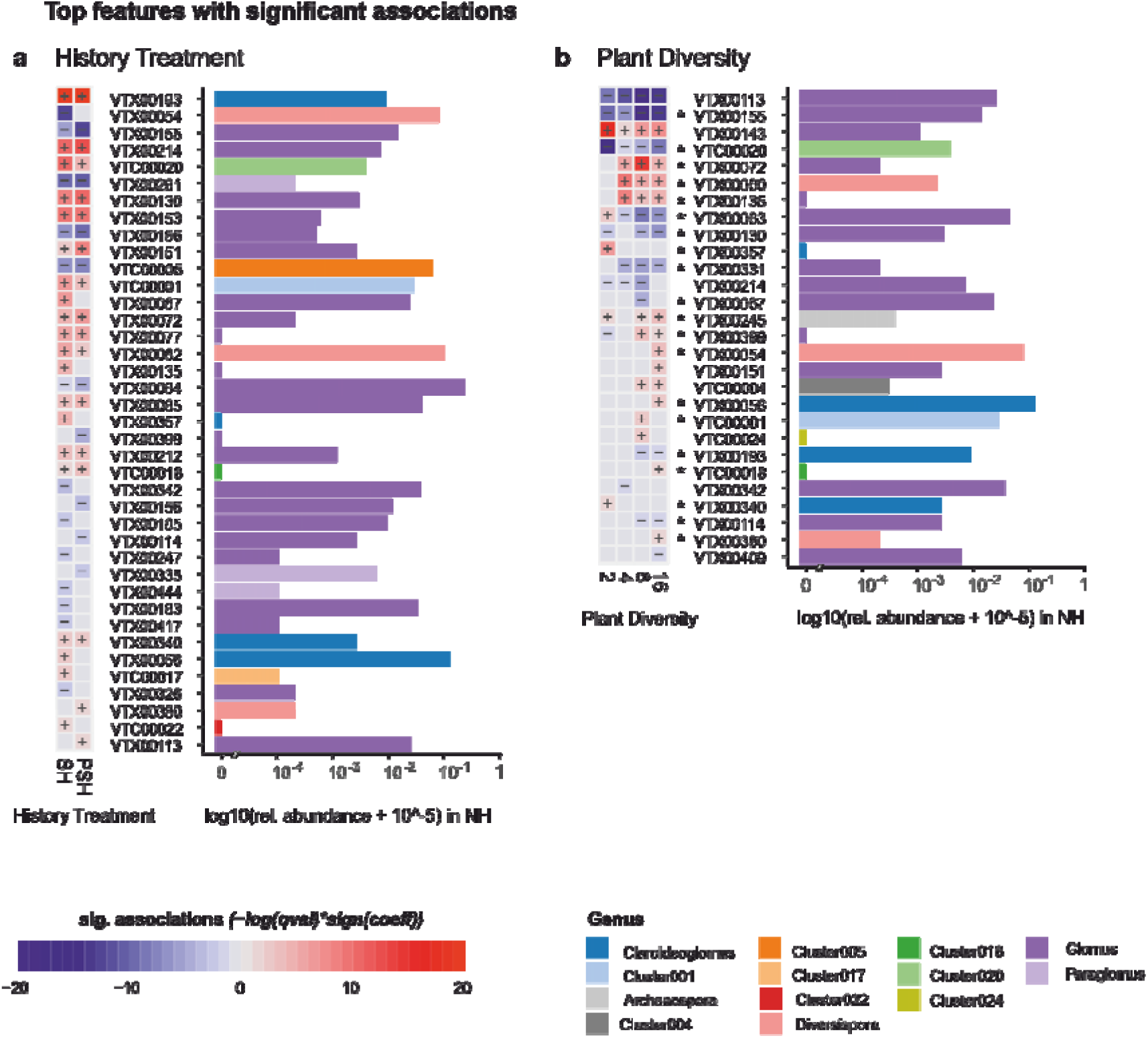
Significantly differential VT showing **(a)** differential abundance in history treatments SH or PSH in comparison to the NH split plots and **(b)** differential abundances of the more diverse plant communities compared to monocultures (factorial comparison) with * indicating significant differences in the linear model. Bar plots indicate relative abundance of VT in NH as base abundance for the differential abundance analysis and colours indicate genus. We found no significant interactions of history treatment and plant diversity. Significance is based on Maaslin2 ZINB model.

Across the plant diversity gradient (**Fig. 4b**), two *Glomus* taxa (VTX00113, VTX00155) were significantly less abundant in the more diverse plots than in monocultures. However, other *Glomus* (VTX00143, VTX00072, VTX00135) and a *Diversispora* (VTX00060) were significantly more abundant at higher plant diversity. While other VT did not show clear trends along the diversity gradient, some VT were only significantly more abundant in the higher diverse plots with 8 or 16 plant species (*Glomus* VTX00399 and VTX00151; *Diversispora* VTX00054; *Claroideoglomus* VTX00056 and *Cluster 4* (*Archaespora*)).

In summary, a total of 39 VT showed significant differential abundance in response to the history treatments and 28 VT responded to the plant diversity gradient. Of these, 20 VT were responding to both the history treatment and the plant diversity, however we could not find an interactive effect of these two factors on differential abundances.

### Plant diversity and history effects on AMF -diversity and turnover

Next, we analysed whether different components of □-diversity of the AMF communities also responded to history or plant diversity (H2). All calculated □-diversity indices were significantly affected by the block in which the plots were located. When adjusting for block effects, VT richness was not different between the 3 history treatments (F = 1.00, p = 0.37, Online Resource 1: **Fig. S2**), but was significantly positively affected by plant diversity (F = 21.02, p < 0.0001). The effect size of plant diversity on AMF richness differed between the treatments, being highest in PSH and lowest in SH (Online Resource 1: **Table S3**). Pielou’s evenness of AMF communities, on the other hand, was affected by the history treatments only (Online Resource 1: **Table S3**). Pielous’s evenness of PSH was significantly higher than NH (t = -3.31, p = 0.001) and SH (t = -2.57, p = 0.012). The history treatments and plant diversity had no significant interactive effect on any of the □-diversity indices.

Species turnover was higher from NH to SH than from SH to PSH (Online Resource 1: **Fig. S3**), suggesting again that soil history has a bigger impact on AMF presences than the plant history. Appearances of species from NH to SH amounted to 40.8 % and to 35.4 % from SH to PSH. The disappearance was higher from NH to SH with 39.3 % than 36.5 % from SH to PSH. In addition to history (F = 21.59, p < 0.001), plant diversity (F = 4.12, p = 0.003) had a significant effect on species turnover, with the turnover being higher in monocultures than mixtures.

### Specificity of AMF communities and effects of plant functional groups

We hypothesised (H3a) that AMF communities specialise over time. To address this, we calculated the φ-coefficient for the frequency of AMF VT co-occurring with specific grassland plant species as measure of specificity per history treatment (**Fig. 5**). Many VT had positive plant species specificities, but the mode of φ was in the slightly negative range. Comparing φ-values against two null-models which preserve VT frequencies or VT frequencies and VT richness per sample, we found only few significant specialisations of AMF for plant species: In NH split plots VTX00156 (*Glomus*) co-occurred with *Gallium mollugo* agg., in SH split plots we found VTX00245 (*Archaespora*) with *Poa pratense* and in PSH split plots no VT was specialised on any plant species (**Table S4**).

**Fig. 5.**
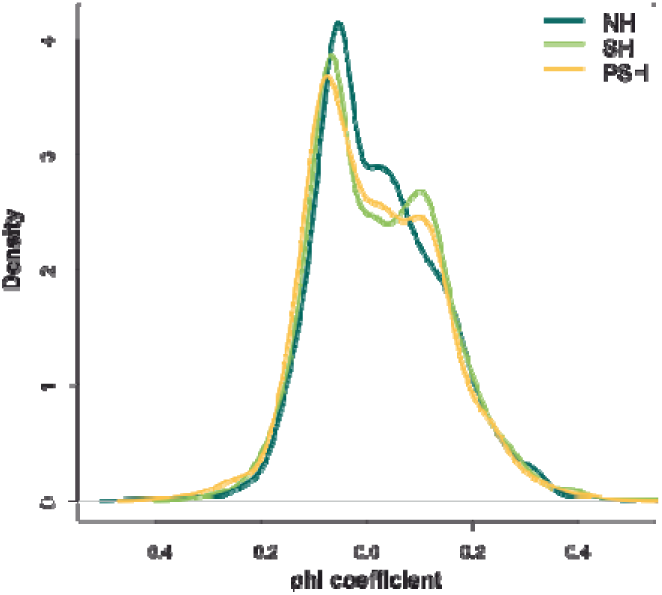
Density of *φ*-coefficients for specificity of each AMF VT to individual plant species with *φ* = -1 being non-specific and *φ* = 1 being highly specific.

### The potential mediating role of AMF communities for plant productivity

Lastly, we tested whether AMF communities may explain ecosystem functioning, given the soil properties and treatments (H4). Significant response of aboveground plant biomass, microbial biomass, microbial respiration and other factors to the plant and soil history treatments had been previously shown in Vogel et al (2019). In accordance with those findings, in 2021 aboveground plant biomass was primarily determined by plant diversity (F = 28.47, p < 0.001) but also by the history treatment (F = 10.74, p < 0.001), with the plant diversity slope being steepest in PSH split-plots (Online Resource 1: **Fig. S4** and **Table S5**), but both plant diversity and history effects were weaker at this later time point. Plant biomass of sown species was significantly higher with legumes (t = 5.52, p < 0.01) or grasses (t = 3.80, p < 0.01) present and also at higher functional plant diversity (F = 25.70, p < 0.01). Unlike the aboveground plant biomass, root biomass was not affected by history treatment (F =1.39, p = 0.25) but by plant diversity (F = 15.72, p < 0.001) and presence of legumes (F = 5.37, p = 0.02) and grasses (F = 5.03, p = 0.03). We found no interaction of history treatment and plant diversity on plant biomass.

To understand how much of the variation in plant and microbial biomass can be explained through influence of the AMF communities, we performed mediation analysis (Online Resource 1: **Table S6**). Only 0.84 % of variation in soil P (p = 0.02), but 2.69 % variation in SWC (p < 0.001) was potentially mediated by AMF. We found no evidence for significant mediation of history treatment and plant diversity effects on aboveground plant community biomass by AMF communities (p = 0.24), but 1.45 % of variations in aboveground legume biomass was potentially mediated through AMF communities. None of the variation in grass (p = 0.26) or herb (p = 0.69) aboveground biomass introduced by history treatment, plant diversity and soil P and SWC were likely mediated through AMF. Belowground, differentiating between coarse and fine root biomass, we found potential mediation effects of AMF on 1.02 % variation of fine root biomass (p < 0.01). To summarise, the potential for the overall bulk soil AMF community composition to be the mediator of the plant diversity and history effects was found to be limited.

## Discussion

Here, we present results of the ΔBEF Experiment (Vogel *et al*., 2019) to test community age effects as well as plant community-specific soil history effects on AMF communities in grasslands. Our analyses show that AMF communities assemble over time and are shaped by plant diversity, but we find no strengthening of plant-AMF diversity relationships over time and limited evidence for the potential of bulk soil AMF communities as lone drivers of BEF relationships.

### Plant history effects are transient while lack of common soil history has long-lasting effects

The experimental plant diversity gradient has previously been established as a factor shaping AMF communities in bulk soil of the Jena Experiment (König *et al*., 2010a; Dassen *et al*., 2017). We expected to see recapitulation of plant diversity effects with re-establishment of the experimental grasslands and indeed observed a plant diversity effect on relative abundances of VT and overall AMF community composition in all history treatments. In addition, the identity of the present plant functional groups played a role, as previously described (König *et al*., 2010a; Dassen *et al*., 2017; Maciá-Vicente *et al*., 2023). We hypothesised common soil history would lead to strong shaping of AMF communities (H1a), while common plant history would additionally drive AMF community composition (H1b). Our analyses showed that the plant diversity effect increased with plant history: the plant diversity effect was stronger in split plots with plant community history (PSH) than in the other split plots. Plus, divergence of AMF communities with soil history was more pronounced in more diverse plant communities, suggesting convergence or more stability of soil processes at higher diversity.

However, within a plot of the same plant community, AMF communities in NH split plots were equally dissimilar from those in split plots with just soil history (SH) as those from split plots with both plant and soil history (PSH). This suggests that soil history was more important for overall AMF community assemblage and additional plant history had less of an impact. Comparing our results after 5 years of re-establishment with those of Macía-Vicente et al (2023) who compared SH and PSH in the same experiment after 1 year of re-establishment, we see that while plant community and diversity effects did drive AMF community composition after 1 year, these effects seems to have disappeared with time. We did not find significant effects of plant history anymore, indicating potentially saturating effects of plant community age between 1 and 5 years. This would be consistent with the observations that, while plants have been shown to drive AMF colonisation of soil patches (in ‘t Zandt *et al*., 2022), plant diversity effects on many soil processes and communities establish after a time lag of ∼ 4 years (Eisenhauer *et al*., 2010; Eisenhauer, 2012). On the other hand, Macía-Vicente et al (2023) analysed root-associated fungi and the difference in plant history effects may be due to the larger influence of the plant community on those than on bulk soil. In bulk soil, it seems that soil legacy effects take more time to develop than plant community effects, as differences in AMF communities and turnover between NH and SH were still present 5 years after establishment, when the effect of plant history had already vanished. The experiment was set up in a formerly arable field and, due to discontinuation of fertilisation, showed changes in nutrient availability over time (Oelmann *et al*., 2007, 2011), which likely continually alters mycorrhizal symbiosis and fungal growth (Antunes *et al*., 2012; Xiao *et al*., 2023) with a visible long-term impact. To disentangle these plant and soil history dynamics further, a fourth treatment of common plant history without soil is considered for the design of a follow-up mesocosm experiment.

Even though we asked questions about fundamental patterns and processes, this apparent difference of temporal effects on plant and soil community assembly can be taken into consideration for more applied approaches e.g. restoration and agriculture where mycorrhizal inoculants have gained interest (Baar, 2008; Rouphael *et al*., 2015; Koziol and Bever, 2017). Even if mycorrhizal inoculants have been shown to immediately increase plant productivity (Hoeksema *et al*., 2010; Bi, Zhang and Zou, 2018), such approaches might not have the desired positive long-term effects, if soil legacies shift or suppress AMF community assembly or alternatively take years to reach full potential (Wubs *et al*., 2019).

### AM fungal diversity and plant diversity relationship lasts over time

We further tried to answer whether plant diversity positively affects AMF diversity in the different experimental grassland ages and whether this relationship is stronger or weaker with increasing community age (H2). Contrary to early findings by Dassen et al (2017), we did find a positive effect of plant diversity on AMF diversity in the Jena Experiment field site. In our analysis, this relationship remained but did not strengthen with increased common history.

However, this only held true for AMF VT richness and Shannon diversity, while AMF Simpson diversity, on which VT relative abundance has a stronger effect than on Shannon diversity, and Pielou’s evenness of AMF communities never increased with plant diversity. Evenness and evenness-affected diversity measures increased with common soil and plant history. Lower evenness in the newly established grassland split plots could be the result of soil disturbance and new plants having increased nutrient needs for initial growth of the plant communities (Jasper, Abbott and Robson, 1991; Hart and Reader, 2004; Trejo, Barois and Sangabriel-Conde, 2016; Vogel *et al*., 2019). AMF community structures are shaped not only by plant diversity, but by dynamic abiotic and biotic factors (Ji and Bever, 2016; He *et al*., 2023; Mansfield *et al*., 2023) which counteract domination of few AMF and lead to more even AMF community composition over time. We conclude that AMF diversity is governed by multiple factors, as AMF communities are subject to plant diversity driving AMF richness while community age drives AMF evenness.

### AMF communities do not become more specialised over time

We expected to see a temporal shift of AMF towards communities with more specialists for plant partners and a changing role in e.g., resource partitioning, with ecosystem age (H3). However, we found that the specificity of AMF was low in all plant communities, independent of history treatments. This result might be related to the sampled bulk soil, which was homogenised to represent the whole split plots. Bulk soil, though reflecting the available species pool and containing the highest AMF diversity (Hempel, Renker and Buscot, 2007), might be less informative on specificity of AMF than what could be found when analysing AMF of the roots or the rhizosphere soil (Ramana *et al*., 2023), where interactions of AMF and plants are higher. The results of Macía-Vicente et al (2023) on root fungi were in line with predictions of Buscot (2015) hypothesising older plant communities to favour a smaller species pool of more generalist fungi. The difference could be the result of soil disturbance or input of plant material with increased nutrient needs for initial growth of the plant communities (Jasper, Abbott and Robson, 1991; Hart and Reader, 2004; Trejo, Barois and Sangabriel-Conde, 2016; Vogel *et al*., 2019). Assuming fungi in bulk soil behave similar to those in roots, after 5 years the initial transition may be over and the selection of a smaller more generalist pool of fungi has been completed, which would explain why we could not find differences between the different plant community ages. Concluding, AMF communities do not gain specialists, but the observed turnover of species showed that both plant diversity and common histories affect AMF assemblage.

### Are AMF drivers of biodiversity and ecosystem functioning relationships?

We described how AM fungal communities in grasslands change with time and plant diversity. Since AMFs have been repeatedly proven to enhance plant productivity and resistance (van der Heijden *et al*., 1998; Hoeksema *et al*., 2010; Schnitzer *et al*., 2011; Wagg *et al*., 2011; Koziol and Bever, 2017; Bi, Zhang and Zou, 2018; Allsup, George and Lankau, 2023), but can also stifle productivity at high diversity levels (Klironomos *et al*., 2000), a central question with regards to biodiversity and ecosystem functioning relationships and its dynamics is, whether AMF are drivers of such relationships (H4). For this to be true, we would expect AMF community composition to plant diversity relationships to change according to shared history. However, across all analyses, the interaction term of history treatment and plant diversity was either not significant or explained less than 1 % of the variance. We further showed a limited potential for AMF to mediate plant diversity and history induced variation of plant productivity both above- and belowground.

Can we use any of the results to predict how AMF communities will change in future with more biodiversity loss and increasing soil/plant history and the feedback of these changes to functioning? Plant diversity is easily manipulatable, and soil remains a complex, more unpredictable system. Nevertheless, the present field experiment gives us insights into long-term developments and a basis to look more deeply into plant-soil-microbes interplay shaping an ecosystem and its potential in practical application.

## Statements and Declarations

### Competing interests

A.H.-B. is editorial board member of Biology and Fertility of Soils. The authors have no competing interests to declare that are relevant to the content of this article.

### Author contributions

F.B., N.E., A.W. and A.H.-B. designed the study. C.A., J.H., M.D.S., L.B., and A.W. carried out the soil sampling. M.D.S. performed DNA extraction. C.A. and J.H. prepared and conducted the sequencing. C.A. and A.H.-B. processed the sequence data and C.A. performed the data analysis with support from G.R.vd.P. and A.H.-B. All authors contributed to the interpretation of the data. The manuscript was written by C.A. and A.H.-B. and all authors commented on the manuscript.

### Data availability

The Illumina sequencing data generated in this study is available in the NCBI Sequence Read Archive (BioProject: PRJNA988299). Further data on biomasses and soil parameters is available for requests through the online data repository of The Jena Experiment (JEXIS: https://jexis.uni-jena.de/) under the following dataset IDs: 315, 319, 240, 318, 321, 379, 303, 304. This data, however, by default falls under the Jena Experiment’ data and publication policy and underlies an embargo period of three years from the end of data collection/data assembly to give data owners and collectors time to perform their analysis. These datasets will be made publicly available via JEXIS at a later point of time. The full R script is uploaded to https://github.com/cyn-alb/deltaBEF_AMF.

## Acknowledgements

The Jena Experiment, C.A., M.D.S, and L.B. are funded by the DFG (FOR 5000). We thank the technical staff of the Jena Experiment for their work in maintaining the experimental field site and the many student helpers for weeding of the experimental plots and support during measurements. We further thank Akanksha Rai, Markus Lange, and Aaron Fox who were heavily involved in taking and processing samples. We thank Beatrix Schnabel for technical support with sample preparation and sequencing. Data processing was performed at the High-Performance Computing (HPC) Cluster EVE, a joint effort of the Helmholtz Centre for Environmental Research–UFZ and the German Centre for Integrative Biodiversity Research (iDiv) Halle-Jena-Leipzig, and we thank Christian Krause and the other administrators for excellent support. We further want to thank Gerlinde Kratzsch, Angelos Amyntas, Annette Jesch, Harald Neidhardt, Alfred Lochner and Yuanyuan Huang, who generated and provided data of the ΔBEF experiment used in this study. N.E. acknowledges support of iDiv funded by the German Research Foundation (DFG – FZT 118, 202548816) and Ei 862/29-1. A.W. acknowledges support of iDiv Flexpool funding and WE 3556/7-1.

## Notes

https://jexis.uni-jena.de

### Competing Interest Statement

The authors have declared no competing interest.

## References

1. Albracht, C. et al. (2023) ‘Effects of recurrent summer droughts on arbuscular mycorrhizal and total fungal communities in experimental grasslands differing in plant diversity and community composition’, Frontiers in Soil Science, 3. Available at: https://www.frontiersin.org/articles/10.3389/fsoil.2023.1129845 (Accessed: 26 June 2023).

2. Allsup, C.M., George, I. and Lankau, R.A. (2023) ‘Shifting microbial communities can enhance tree tolerance to changing climates’, Science, 380(6647), pp. 835–840. Available at: 10.1126/science.adf2027.

3. Antoninka, A., Reich, P.B. and Johnson, N.C. (2011) ‘Seven years of carbon dioxide enrichment, nitrogen fertilization and plant diversity influence arbuscular mycorrhizal fungi in a grassland ecosystem’, New Phytologist, 192(1), pp. 200–214. Available at: 10.1111/j.1469-8137.2011.03776.x.

4. Antunes, P.M. et al. (2012) ‘Long-term effects of soil nutrient deficiency on arbuscular mycorrhizal communities’, Functional Ecology, 26(2), pp. 532–540. Available at: 10.1111/j.1365-2435.2011.01953.x.

5. Baar, J. (2008) ‘From Production to Application of Arbuscular Mycorrhizal Fungi in Agricultural Systems: Requirements and Needs’, in A. Varma (ed.) Mycorrhiza: State of the Art, Genetics and Molecular Biology, Eco-Function, Biotechnology, Eco-Physiology, Structure and Systematics . Berlin, Heidelberg: Springer, pp. 361–373. Available at: 10.1007/978-3-540-78826-3_18.

6. Barry, K.E. et al. (2018) ‘The Future of Complementarity: Disentangling Causes from Consequences’, Trends in Ecology & Evolution, 34(2), pp. 167–180. Available at: 10.1016/j.tree.2018.10.013.

7. Barry, K.E. et al. (2019) ‘Limited evidence for spatial resource partitioning across temperate grassland biodiversity experiments’, Ecology, 101(1), p. e02905. Available at: 10.1002/ecy.2905.

8. Berger, F. and Gutjahr, C. (2021) ‘Factors affecting plant responsiveness to arbuscular mycorrhiza’, Current Opinion in Plant Biology, 59, p. 101994. Available at: 10.1016/j.pbi.2020.101994.

9. Bi, Y., Zhang, Y. and Zou, H. (2018) ‘Plant growth and their root development after inoculation of arbuscular mycorrhizal fungi in coal mine subsided areas’, International Journal of Coal Science & Technology, 5(1), pp. 47–53. Available at: 10.1007/s40789-018-0201-x.

10. Bro, R. (1997) ‘PARAFAC. Tutorial and applications’, Chemometrics and Intelligent Laboratory Systems [Preprint].

11. Bro, R. (2023) ‘The N-Way Toolbox’, *MATLAB Central File Exchange* [Preprint]. Available at: https://www.mathworks.com/matlabcentral/fileexchange/1088-the-n-way-toolbox.

12. Bro, R. and Smilde, A.K. (2003) ‘Centering and scaling in component analysis’, Journal of Chemometrics, 17(1), pp. 16–33. Available at: 10.1002/cem.773.

13. Burrows, R.L. and Pfleger, F.L. (2002) ‘Arbuscular mycorrhizal fungi respond to increasing plant diversity’, Canadian Journal of Botany, 80(2), pp. 120–130. Available at: 10.1139/b01-138.

14. Buscot, F. (2015) ‘Implication of evolution and diversity in arbuscular and ectomycorrhizal symbioses’, Journal of Plant Physiology, 172, pp. 55–61. Available at: 10.1016/j.jplph.2014.08.013.

15. Callahan, B.J. et al. (2016) ‘DADA2: High-resolution sample inference from Illumina amplicon data’, Nature Methods, 13(7), pp. 581–583. Available at: 10.1038/nmeth.3869.

16. Cardinale, B.J. et al. (2007) ‘Impacts of plant diversity on biomass production increase through time because of species complementarity’, Proceedings of the National Academy of Sciences, 104(46), pp. 18123–18128. Available at: 10.1073/pnas.0709069104.

17. Cardinale, B.J. (2011) ‘Biodiversity improves water quality through niche partitioning’, Nature, 472(7341), pp. 86–89. Available at: 10.1038/nature09904.

18. Carroll, J.D. and Chang, J.-J. (1970) ‘Analysis of individual differences in multidimensional scaling via an n-way generalization of “Eckart-Young” decomposition’, Psychometrika, 35(3), pp. 283–319. Available at: 10.1007/BF02310791.

19. Chowdhury, S. et al. (2022) ‘Plants with arbuscular mycorrhizal fungi efficiently acquire Nitrogen from substrate additions by shaping the decomposer community composition and their net plant carbon demand’, Plant and Soil, 475(1), pp. 473–490. Available at: 10.1007/s11104-022-05380-x.

20. Chytrý, M. et al. (2002) ‘Determination of diagnostic species with statistical fidelity measures’, Journal of Vegetation Science, 13(1), pp. 79–90. Available at: 10.1111/j.1654-1103.2002.tb02025.x.

21. Dassen, S. et al. (2017) ‘Differential responses of soil bacteria, fungi, archaea and protists to plant species richness and plant functional group identity’, Molecular Ecology, 26(15), pp. 4085–4098. Available at: 10.1111/mec.14175.

22. Eisenhauer, N. et al. (2010) ‘Plant diversity effects on soil microorganisms support the singular hypothesis’, Ecology, 91(2), pp. 485–496. Available at: 10.1890/08-2338.1.

23. Eisenhauer, N. (2012) ‘Aboveground–belowground interactions as a source of complementarity effects in biodiversity experiments’, Plant and Soil, 351(1), pp. 1–22. Available at: 10.1007/s11104-011-1027-0.

24. Eisenhauer, N. et al. (2017) ‘Root biomass and exudates link plant diversity with soil bacterial and fungal biomass’, Scientific Reports, 7(1), p. 44641. Available at: 10.1038/srep44641.

25. Eisenhauer, N., et al. (2019) ‘Chapter One - A multitrophic perspective on biodiversity–ecosystem functioning research’, in N. Eisenhauer, D.A. Bohan, and A.J. Dumbrell (eds) Advances in Ecological Research. Academic Press (Mechanisms underlying the relationship between biodiversity and ecosystem function), pp. 1–54. Available at: 10.1016/bs.aecr.2019.06.001.

26. Fargione, J. et al. (2007) ‘From selection to complementarity: shifts in the causes of biodiversity– productivity relationships in a long-term biodiversity experiment’, Proceedings of the Royal Society B: Biological Sciences, 274(1611), pp. 871–876. Available at: 10.1098/rspb.2006.0351.

27. Fornara, D.A. and Tilman, D. (2008) ‘Plant functional composition influences rates of soil carbon and nitrogen accumulation’, Journal of Ecology, 96(2), pp. 314–322. Available at: 10.1111/j.1365-2745.2007.01345.x.

28. García-Valdés, R., Bugmann, H. and Morin, X. (2018) ‘Climate change-driven extinctions of tree species affect forest functioning more than random extinctions’, Diversity and Distributions, 24(7), pp. 906–918. Available at: 10.1111/ddi.12744.

29. Guzman, A. et al. (2021) ‘Crop diversity enriches arbuscular mycorrhizal fungal communities in an intensive agricultural landscape’, New Phytologist, 231(1), pp. 447–459. Available at: 10.1111/nph.17306.

30. Habekost, M. et al. (2008) ‘Seasonal changes in the soil microbial community in a grassland plant diversity gradient four years after establishment’, Soil Biology and Biochemistry, 40(10), pp. 2588– 2595. Available at: 10.1016/j.soilbio.2008.06.019.

31. Hallett, L.M. et al. (2016) ‘codyn: An r package of community dynamics metrics’, Methods in Ecology and Evolution, 7(10), pp. 1146–1151. Available at: 10.1111/2041-210X.12569.

32. Harshman, R.A. (1970) ‘Foundations of the PARAFAC procedure: Models and conditions for an “explanatory” multimodal factor analysis’.

33. Hart, M. and Reader, I. (2004) ‘Do arbuscular mycorrhizal fungi recover from disturbance differently?’, Trop. Ecol., 45.

34. He, X. et al. (2023) ‘Accuracy of mutual predictions of plant and microbial communities vary along a successional gradient in an alpine glacier forefield’, Frontiers in Plant Science, 13, p. 1017847. Available at: 10.3389/fpls.2022.1017847.

35. Hector, A. et al. (1999) ‘Plant Diversity and Productivity Experiments in European Grasslands’, Science, 286(5442), pp. 1123–1127. Available at: 10.1126/science.286.5442.1123.

36. van der Heijden, M.G.A. et al. (1998) ‘Mycorrhizal fungal diversity determines plant biodiversity, ecosystem variability and productivity’, Nature, 396(6706), pp. 69–72. Available at: 10.1038/23932.

37. Hempel, S., Renker, C. and Buscot, F. (2007) ‘Differences in the species composition of arbuscular mycorrhizal fungi in spore, root and soil communities in a grassland ecosystem’, Environmental Microbiology, 9(8), pp. 1930–1938. Available at: 10.1111/j.1462-2920.2007.01309.x.

38. Hoeksema, J.D. et al. (2010) ‘A meta-analysis of context-dependency in plant response to inoculation with mycorrhizal fungi’, Ecology Letters, 13(3), pp. 394–407. Available at: 10.1111/j.1461-0248.2009.01430.x.

39. Hong, P. et al. (2022) ‘Biodiversity promotes ecosystem functioning despite environmental change’, Ecology Letters, 25(2), pp. 555–569. Available at: 10.1111/ele.13936.

40. Hu, Y.-J. and Satten, G.A. (2022) LDM: Testing Hypotheses about the Microbiome using the Linear Decomposition Model, Version 5.0. Available at: https://github.com/yijuanhu/LDM.

41. Jasper, D.A., Abbott, L.K. and Robson, A.D. (1991) ‘The effect of soil disturbance on vesicular— arbuscular mycorrhizal fungi in soils from different vegetation types’, New Phytologist, 118(3), pp. 471–476. Available at: 10.1111/j.1469-8137.1991.tb00029.x.

42. Ji, B. and Bever, J.D. (2016) ‘Plant preferential allocation and fungal reward decline with soil phosphorus: implications for mycorrhizal mutualism’, Ecosphere, 7(5), p. e01256. Available at: 10.1002/ecs2.1256.

43. Klironomos, J.N. et al. (2000) ‘The influence of arbuscular mycorrhizae on the relationship between plant diversity and productivity’, Ecology Letters, 3(2), pp. 137–141. Available at: 10.1046/j.1461-0248.2000.00131.x.

44. König, S. et al. (2010a) ‘TaqMan Real-Time PCR Assays To Assess Arbuscular Mycorrhizal Responses to Field Manipulation of Grassland Biodiversity: Effects of Soil Characteristics, Plant Species Richness, and Functional Traits’, Applied and Environmental Microbiology, 76(12), pp. 3765–3775. Available at: 10.1128/AEM.02951-09.

45. König, S. et al. (2010b) ‘TaqMan Real-Time PCR Assays To Assess Arbuscular Mycorrhizal Responses to Field Manipulation of Grassland Biodiversity: Effects of Soil Characteristics, Plant Species Richness, and Functional Traits’, Applied and Environmental Microbiology, 76(12), pp. 3765–3775. Available at: 10.1128/AEM.02951-09.

46. Koziol, L. and Bever, J.D. (2017) ‘The missing link in grassland restoration: arbuscular mycorrhizal fungi inoculation increases plant diversity and accelerates succession’, Journal of Applied Ecology, 54(5), pp. 1301–1309. Available at: 10.1111/1365-2664.12843.

47. Kuyper, T.W. and Jansa, J. (2023) ‘Arbuscular mycorrhiza: advances and retreats in our understanding of the ecological functioning of the mother of all root symbioses’, Plant and Soil [Preprint]. Available at: 10.1007/s11104-023-06045-z.

48. Lange, M. et al. (2014) ‘Biotic and Abiotic Properties Mediating Plant Diversity Effects on Soil Microbial Communities in an Experimental Grassland’, PLOS ONE, 9(5), p. e96182. Available at: 10.1371/journal.pone.0096182.

49. Lange, M. et al. (2015) ‘Plant diversity increases soil microbial activity and soil carbon storage’, Nature Communications, 6(1), p. 6707. Available at: 10.1038/ncomms7707.

50. Lange, M. et al. (2023) ‘Increased soil carbon storage through plant diversity strengthens with time and extends into the subsoil’, Global Change Biology, 29(9), pp. 2627–2639. Available at: 10.1111/gcb.16641.

51. Lee, J., Lee, S. and Young, J.P.W. (2008) ‘Improved PCR primers for the detection and identification of arbuscular mycorrhizal fungi’, FEMS Microbiology Ecology, 65(2), pp. 339–349. Available at: 10.1111/j.1574-6941.2008.00531.x.

52. Maciá-Vicente, J.G. et al. (2023) ‘The structure of root-associated fungal communities is related to the long-term effects of plant diversity on productivity’, Molecular Ecology, n/a(n/a). Available at: 10.1111/mec.16956.

53. Mallick, H. et al. (2021) ‘Multivariable association discovery in population-scale meta-omics studies’, PLOS Computational Biology, 17(11), p. e1009442. Available at: 10.1371/journal.pcbi.1009442.

54. Mansfield, T.M. et al. (2023) ‘Niche differentiation of Mucoromycotinian and Glomeromycotinian arbuscular mycorrhizal fungi along a 2-million-year soil chronosequence’, Mycorrhiza [Preprint]. Available at: 10.1007/s00572-023-01111-x.

55. McMurdie, P.J. and Holmes, S. (2013) ‘phyloseq: An R Package for Reproducible Interactive Analysis and Graphics of Microbiome Census Data’, PLOS ONE, 8(4), p. e61217. Available at: 10.1371/journal.pone.0061217.

56. Morgan, B.S.T. and Egerton-Warburton, L.M. (2017) ‘Barcoded NS31/AML2 primers for sequencing of arbuscular mycorrhizal communities in environmental samples’, Applications in Plant Sciences, 5(8), p. 1700017. Available at: 10.3732/apps.1700017.

57. Murphy, J. and Riley, J.P. (1962) ‘A modified single solution method for the determination of phosphate in natural waters’, Analytica chimica acta, 27, pp. 31–36.

58. Oelmann, Y. et al. (2007) ‘Soil and plant nitrogen pools as related to plant diversity in an experimental grassland’, Soil Science Society of America Journal, 71(3), pp. 720–729.

59. Oelmann, Y. et al. (2011) ‘Plant diversity effects on aboveground and belowground N pools in temperate grassland ecosystems: development in the first 5 years after establishment’, Global Biogeochemical Cycles, 25(2).

60. Oksanen, J. et al. (2019) ‘Package “vegan”’, Community ecology package, version, 2(9).

61. Olsen, S.R. (1954) Estimation of available phosphorus in soils by extraction with sodium bicarbonate . US Department of Agriculture.

62. Öpik, M. et al. (2010) ‘The online database MaarjAM reveals global and ecosystemic distribution patterns in arbuscular mycorrhizal fungi (Glomeromycota)’, New Phytologist, 188(1), pp. 223–241. Available at: 10.1111/j.1469-8137.2010.03334.x.

63. Price, M.N., Dehal, P.S. and Arkin, A.P. (2010) ‘FastTree 2 – Approximately Maximum-Likelihood Trees for Large Alignments’, PLOS ONE, 5(3), p. e9490. Available at: 10.1371/journal.pone.0009490.

64. Quast, C. et al. (2013) ‘The SILVA ribosomal RNA gene database project: improved data processing and web-based tools’, Nucleic Acids Research, 41(D1), pp. D590–D596. Available at: 10.1093/nar/gks1219.

65. R Core Team (2017) ‘R: A language and environment for statistical computing’. R Foundation for Statistical Computing, Vienna, Austria. Available at: https://www.R-project.org/.

66. Ramana, J.V. et al. (2023) ‘Root diameter, host specificity and arbuscular mycorrhizal fungal community composition among native and exotic plant species’, New Phytologist, n/a(n/a). Available at: 10.1111/nph.18911.

67. Reich, P.B. et al. (2012) ‘Impacts of Biodiversity Loss Escalate Through Time as Redundancy Fades’, Science, 336(6081), pp. 589–592. Available at: 10.1126/science.1217909.

68. Roscher, C. et al. (2004) ‘The role of biodiversity for element cycling and trophic interactions: an experimental approach in a grassland community’, Basic and Applied Ecology, 5(2), pp. 107–121. Available at: 10.1078/1439-1791-00216.

69. Roscher, C. et al. (2012) ‘Using Plant Functional Traits to Explain Diversity–Productivity Relationships’, PLOS ONE, 7(5), p. e36760. Available at: 10.1371/journal.pone.0036760.

70. Rouphael, Y. et al. (2015) ‘Arbuscular mycorrhizal fungi act as biostimulants in horticultural crops’, Scientia Horticulturae, 196, pp. 91–108. Available at: 10.1016/j.scienta.2015.09.002.

71. Rudgers, J.A. et al. (2020) ‘Climate Disruption of Plant-Microbe Interactions’, Annual Review of Ecology, Evolution, and Systematics, 51(1), pp. 561–586. Available at: 10.1146/annurev-ecolsys-011720-090819.

72. Schloss, P.D. et al. (2009) ‘Introducing mothur: Open-Source, Platform-Independent, Community-Supported Software for Describing and Comparing Microbial Communities’, Applied and Environmental Microbiology, 75(23), pp. 7537–7541. Available at: 10.1128/AEM.01541-09.

73. Schmid, M.W. et al. (2021) ‘Effects of plant community history, soil legacy and plant diversity on soil microbial communities’, Journal of Ecology, 109(8), pp. 3007–3023.

74. Schnitzer, S.A. et al. (2011) ‘Soil microbes drive the classic plant diversity–productivity pattern’, Ecology, 92(2), pp. 296–303. Available at: 10.1890/10-0773.1.

75. Simon, L., Lalonde, M. and Bruns, T.D. (1992) ‘Specific amplification of 18S fungal ribosomal genes from vesicular-arbuscular endomycorrhizal fungi colonizing roots’, Applied and Environmental Microbiology, 58(1), pp. 291–295. Available at: 10.1128/aem.58.1.291-295.1992.

76. Smith, S. and Read, D. (2008) Mycorrhizal Symbiosis. Academic Press. Available at: 10.1016/B978-0-12-370526-6.X5001-6.

77. Stamatakis, A. (2014) ‘RAxML version 8: a tool for phylogenetic analysis and post-analysis of large phylogenies’, Bioinformatics, 30(9), pp. 1312–1313. Available at: 10.1093/bioinformatics/btu033.

78. Steudel, B. et al. (2016) ‘Contrasting biodiversity–ecosystem functioning relationships in phylogenetic and functional diversity’, New Phytologist, 212(2), pp. 409–420. Available at: 10.1111/nph.14054.

79. Strecker, T. et al. (2016) ‘Functional composition of plant communities determines the spatial and temporal stability of soil microbial properties in a long-term plant diversity experiment’, Oikos, 125(12), pp. 1743–1754. Available at: 10.1111/oik.03181.

80. in ‘t Zandt, D., et al. (2022) ‘Plant life-history traits rather than soil legacies determine colonisation of soil patches in a multi-species grassland’, Journal of Ecology, 110(4), pp. 889–901. Available at: 10.1111/1365-2745.13850.

81. Templ, M., Hron, K. and Filzmoser, P. (2011) robCompositions: an R-package for robust statistical analysis of compositional data. John Wiley and Sons.

82. Thakur, M.P. et al. (2015) ‘Plant diversity drives soil microbial biomass carbon in grasslands irrespective of global environmental change factors’, Global Change Biology, 21(11), pp. 4076–4085. Available at: 10.1111/gcb.13011.

83. The MathWorks Inc. (2022) ‘MATLAB version 9.14.0.2239454 (R2023a)’, *The MathWorks Inc.* [Preprint]. Available at: https://www.mathworks.com.

84. Tilman, D., Isbell, F. and Cowles, J.M. (2014) ‘Biodiversity and Ecosystem Functioning’, Annual Review of Ecology, Evolution, and Systematics, 45(1), pp. 471–493. Available at: 10.1146/annurev-ecolsys-120213-091917.

85. Tilman, D. and Snell-Rood, E.C. (2014) ‘Diversity breeds complementarity’, Nature, 515(7525), pp. 44–45. Available at: 10.1038/nature13929.

86. Trejo, D., Barois, I. and Sangabriel-Conde, W. (2016) ‘Disturbance and land use effect on functional diversity of the arbuscular mycorrhizal fungi’, Agroforestry Systems, 90(2), pp. 265–279. Available at: 10.1007/s10457-015-9852-4.

87. Vogel, A., et al. (2019) ‘A new experimental approach to test why biodiversity effects strengthen as ecosystems age’, in Advances in Ecological Research. Elsevier, pp. 221–264. Available at: 10.1016/bs.aecr.2019.06.006.

88. Vogel, A., Scherer-Lorenzen, M. and Weigelt, A. (2012) ‘Grassland Resistance and Resilience after Drought Depends on Management Intensity and Species Richness’, PLOS ONE, 7(5), p. e36992. Available at: 10.1371/journal.pone.0036992.

89. Wagg, C. et al. (2011) ‘Belowground biodiversity effects of plant symbionts support aboveground productivity’, Ecology Letters, 14(10), pp. 1001–1009. Available at: 10.1111/j.1461-0248.2011.01666.x.

90. Wagg, C. et al. (2014) ‘Soil biodiversity and soil community composition determine ecosystem multifunctionality’, Proceedings of the National Academy of Sciences of the United States of Americ, a 111(14), pp. 5266–5270. Available at: 10.1073/pnas.1320054111.

91. Wagg, C. et al. (2015) ‘Complementarity in both plant and mycorrhizal fungal communities are not necessarily increased by diversity in the other’, Journal of Ecology, 103(5), pp. 1233–1244. Available at: 10.1111/1365-2745.12452.

92. Wagg, C. et al. (2022) ‘Biodiversity–stability relationships strengthen over time in a long-term grassland experiment’, Nature Communications, 13(1), p. 7752. Available at: 10.1038/s41467-022-35189-2.

93. Wahdan, S.F.M. et al. (2021) ‘Organic agricultural practice enhances arbuscular mycorrhizal symbiosis in correspondence to soil warming and altered precipitation patterns’, Environmental Microbiology, 23(10), pp. 6163–6176. Available at: 10.1111/1462-2920.15492.

94. Wang, C. et al. (2022) ‘Plant diversity has stronger linkage with soil fungal diversity than with bacterial diversity across grasslands of northern China’, Global Ecology and Biogeography, 31(5), pp. 886–900. Available at: 10.1111/geb.13462.

95. Weißbecker, C. et al. (2019) ‘Linking Soil Fungal Generality to Tree Richness in Young Subtropical Chinese Forests’, Microorganisms, 7(11), p. 547. Available at: 10.3390/microorganisms7110547.

96. Weißbecker, C., Schnabel, B. and Heintz-Buschart, A. (2020) ‘Dadasnake, a Snakemake implementation of DADA2 to process amplicon sequencing data for microbial ecology’, GigaScience, 9(12), p. giaa135. Available at: 10.1093/gigascience/giaa135.

97. Weisser, W.W. et al. (2017) ‘Biodiversity effects on ecosystem functioning in a 15-year grassland experiment: Patterns, mechanisms, and open questions’, Basic and Applied Ecology, 23, pp. 1–73. Available at: 10.1016/j.baae.2017.06.002.

98. Wickham, H. (2009) ggplot2: Elegant Graphics for Data Analysi.s New York, NY: Springer. Available at: 10.1007/978-0-387-98141-3.

99. Wright, A.J. et al. (2015) ‘Flooding disturbances increase resource availability and productivity but reduce stability in diverse plant communities’, Nature Communications, 6(1), p. 6092. Available at: 10.1038/ncomms7092.

100. Wright, A.J. et al. (2017) ‘The Overlooked Role of Facilitation in Biodiversity Experiments’, Trends in Ecology & Evolution, 32(5), pp. 383–390. Available at: 10.1016/j.tree.2017.02.011.

101. Wubet, T. et al. (2006) ‘Two threatened coexisting indigenous conifer species in the dry Afromontane forests of Ethiopia are associated with distinct arbuscular mycorrhizal fungal communities’, Canadian Journal of Botany, 84(10), pp. 1617–1627. Available at: 10.1139/b06-121.

102. Wubs, E.R.J. et al. (2019) ‘Single introductions of soil biota and plants generate long-term legacies in soil and plant community assembly’, Ecology Letters, 22(7), pp. 1145–1151. Available at: 10.1111/ele.13271.

103. Xiao, D. et al. (2023) ‘Soil nutrients and vegetation along a karst slope gradient affect arbuscular mycorrhizal fungi colonization of roots rather than bulk soil AMF diversity’, Plant and Soil, pp. 1–16. Available at: 10.1007/s11104-023-06004-8.

104. Zuppinger-Dingley, D. et al. (2014) ‘Selection for niche differentiation in plant communities increases biodiversity effects’, Nature, 515(7525), pp. 108–111. Available at: 10.1038/nature13869.

105. Zuppinger-Dingley, D. et al. (2016) ‘Plant selection and soil legacy enhance long-term biodiversity effects’, Ecology, 97(4), pp. 918–928. Available at: 10.1890/15-0599.1.

