## Supplementary Data for "Common soil history is more important than plant history for arbuscular mycorrhizal community assembly in an experimental grassland diversity gradient"

The following Supporting Information is available for this article:

**Fig. S1** Relative Abundances

**Fig. S2** α-diversity of AMF

**Fig. S3** AMF species turnover

**Fig. S4** Plant Biomass

**Table S1** List of VTX

**Table S2** List of VTC

**Table S3** α-diversity models

**Table S4** φ-coefficient

**Table S5** ANOVA on plant and microbial biomasses

**Table S6** Mediation analyses

**Fig. S1** Average relative abundances of AMF genera per history treatment and plant diversity level.

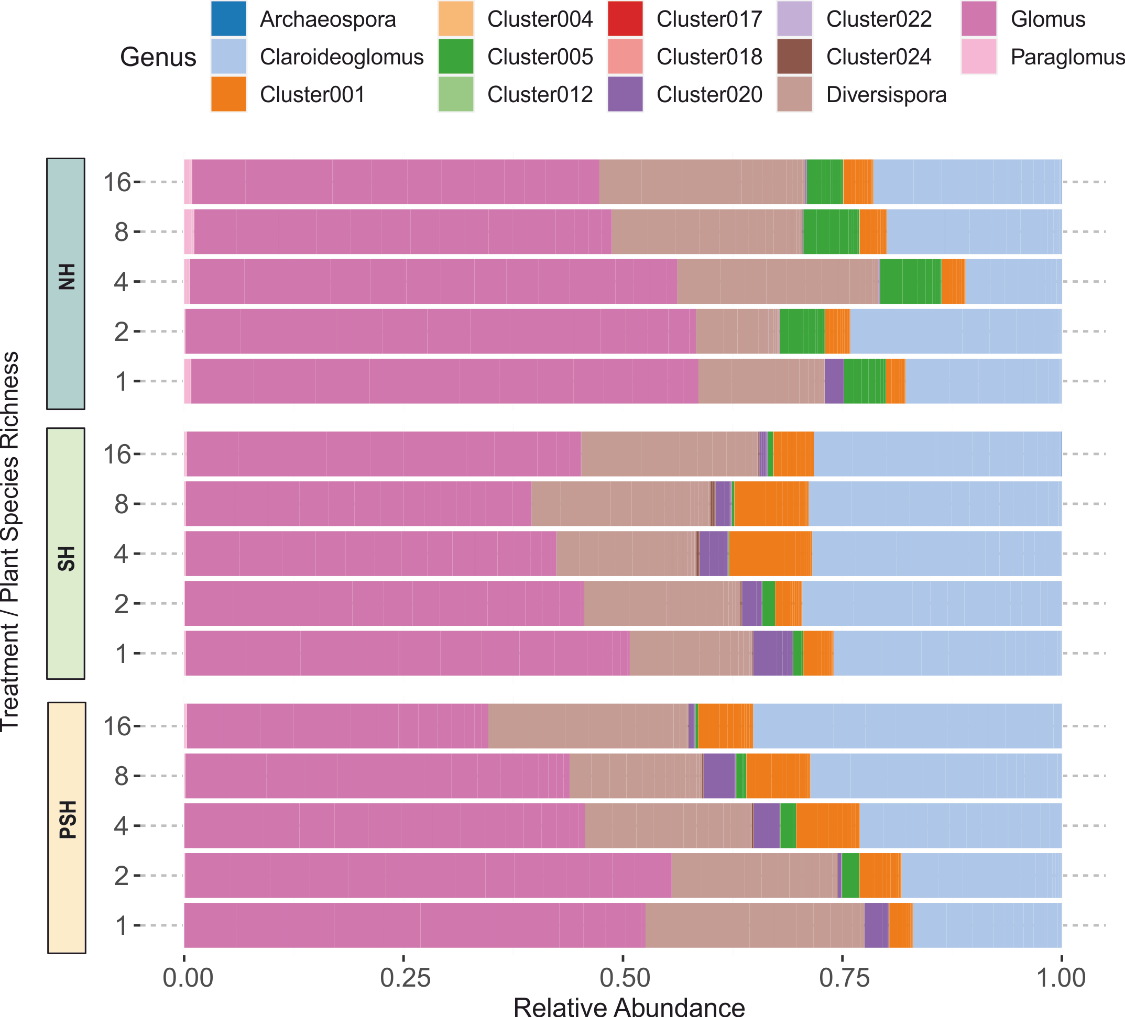

**Fig. S2** Alpha-diversity of AMF communities at the community level calculated as **(a)** VT richness, **(b)** Shannon diversity, **(c)** Simpson diversity and **(d)** Pielou’s evenness indices.

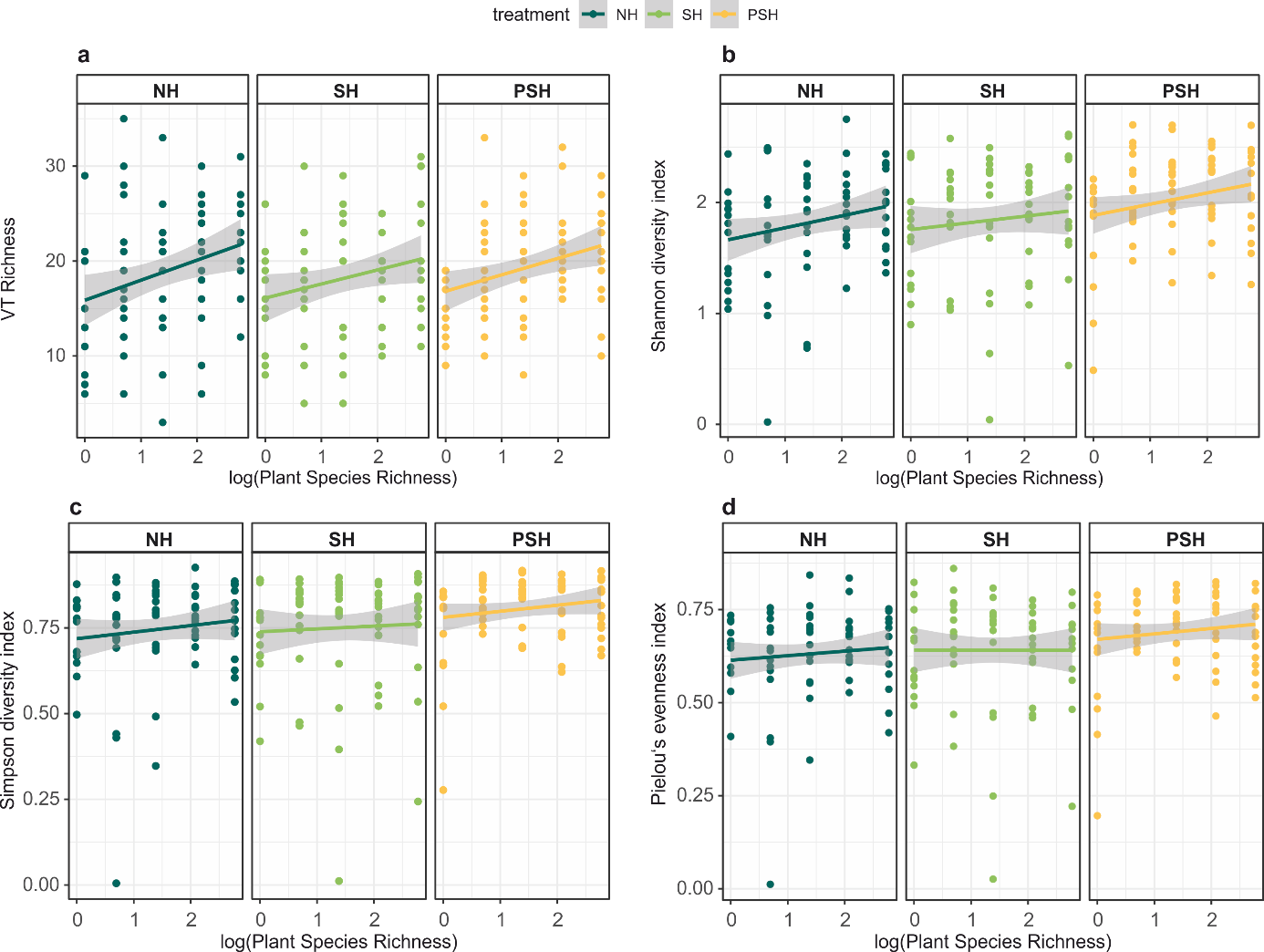

**Fig. S3** Species turnover across plant / soil history shown as appearance, disappearance and total turnover (*(species lost + species gained) / total species*) observed from NH to SH and from SH to PSH within each plant diversity level.

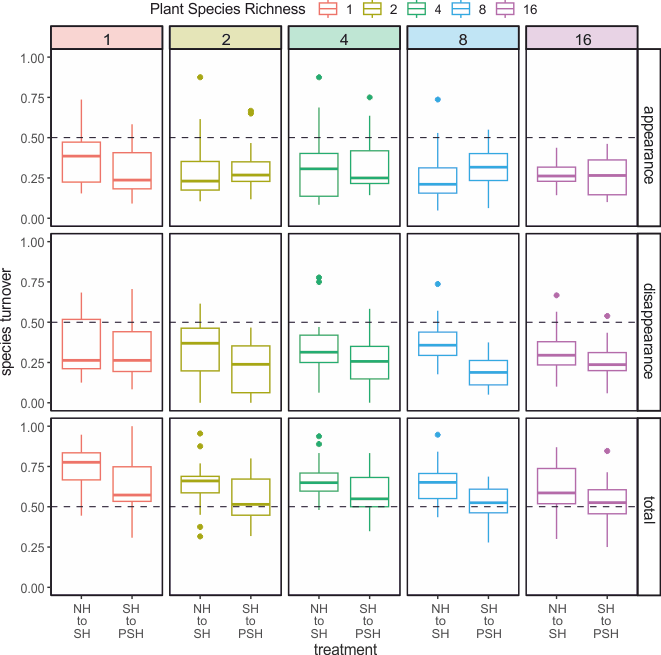

**Fig. S4** Total aboveground plant biomass of target species and biomass per plant functional group (grasses, herbs, legumes) per history treatment and plant diversity.

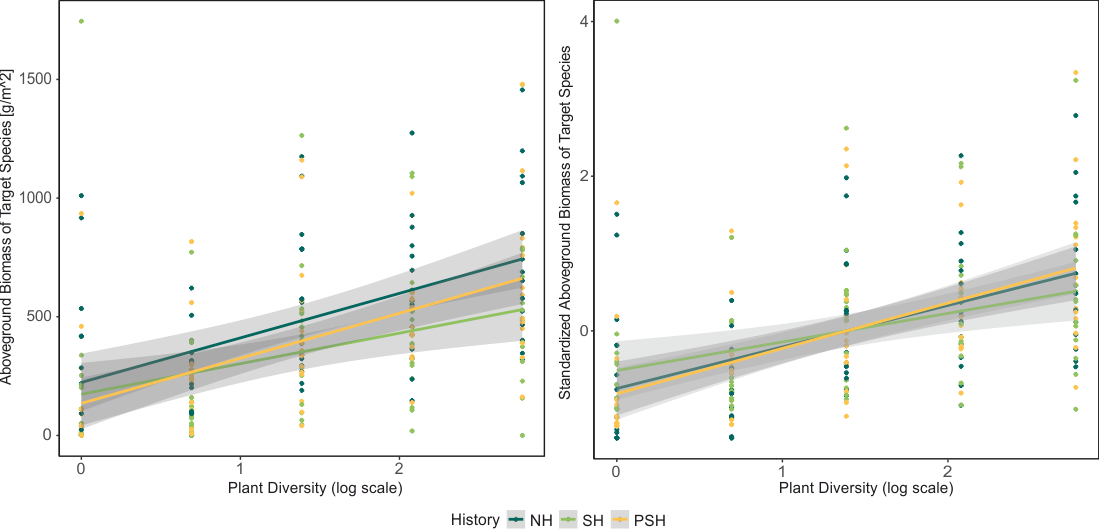

**Table S1** List of VTX and their assigned taxonomy according to MaarjAM database (accessed April 28th 2023).

| **VT** | **Family** | **Genus** | **VT** | **Family** | **Genus** |
| --- | --- | --- | --- | --- | --- |
| **VTX00001** | *Paraglomeraceae* | *Paraglomus* | **VTX00222** | *Glomeraceae* | *Glomus* |
| **VTX00008** | *Archaeosporaceae* | *Archaeospora* | **VTX00225** | *Claroideoglomeraceae* | *Claroideoglomus* |
| **VTX00026** | *Acaulosporaceae* | *Acaulospora* | **VTX00237** | *Claroideoglomeraceae* | *Claroideoglomus* |
| **VTX00052** | *Gigasporaceae* | *Scutellospora* | **VTX00238** | *Paraglomeraceae* | *Paraglomus* |
| **VTX00053** | *Glomeraceae* | *Glomus* | **VTX00242** | *Ambisporaceae* | *Ambispora* |
| **VTX00054** | *Diversisporaceae* | *Diversispora* | **VTX00245** | *Archaeosporaceae* | *Archaeospora* |
| **VTX00056** | *Claroideoglomeraceae* | *Claroideoglomus* | **VTX00247** | *Glomeraceae* | *Glomus* |
| **VTX00057** | *Claroideoglomeraceae* | *Claroideoglomus* | **VTX00263** | *Diversisporaceae* | *Diversispora* |
| **VTX00060** | *Diversisporaceae* | *Diversispora* | **VTX00265** | *Glomeraceae* | *Glomus* |
| **VTX00061** | *Diversisporaceae* | *Diversispora* | **VTX00276** | *Claroideoglomeraceae* | *Claroideoglomus* |
| **VTX00062** | *Diversisporaceae* | *Diversispora* | **VTX00278** | *Claroideoglomeraceae* | *Claroideoglomus* |
| **VTX00063** | *Glomeraceae* | *Glomus* | **VTX00281** | *Paraglomeraceae* | *Paraglomus* |
| **VTX00064** | *Glomeraceae* | *Glomus* | **VTX00283** | *Ambisporaceae* | *Ambispora* |
| **VTX00065** | *Glomeraceae* | *Glomus* | **VTX00295** | *Glomeraceae* | *Glomus* |
| **VTX00067** | *Glomeraceae* | *Glomus* | **VTX00296** | *Glomeraceae* | *Glomus* |
| **VTX00072** | *Glomeraceae* | *Glomus* | **VTX00297** | *Claroideoglomeraceae* | *Claroideoglomus* |
| **VTX00077** | *Glomeraceae* | *Glomus* | **VTX00301** | *Glomeraceae* | *Glomus* |
| **VTX00085** | *Glomeraceae* | *Glomus* | **VTX00304** | *Glomeraceae* | *Glomus* |
| **VTX00098** | *Glomeraceae* | *Glomus* | **VTX00306** | *Diversisporaceae* | *Diversispora* |
| **VTX00105** | *Glomeraceae* | *Glomus* | **VTX00307** | *Glomeraceae* | *Glomus* |
| **VTX00113** | *Glomeraceae* | *Glomus* | **VTX00308** | *Paraglomeraceae* | *Paraglomus* |
| **VTX00114** | *Glomeraceae* | *Glomus* | **VTX00309** | *Glomeraceae* | *Glomus* |
| **VTX00115** | *Glomeraceae* | *Glomus* | **VTX00312** | *Glomeraceae* | *Glomus* |
| **VTX00118** | *Glomeraceae* | *Glomus* | **VTX00323** | *Glomeraceae* | *Glomus* |
| **VTX00125** | *Glomeraceae* | *Glomus* | **VTX00326** | *Glomeraceae* | *Glomus* |
| **VTX00129** | *Glomeraceae* | *Glomus* | **VTX00331** | *Glomeraceae* | *Glomus* |
| **VTX00130** | *Glomeraceae* | *Glomus* | **VTX00333** | *Glomeraceae* | *Glomus* |
| **VTX00135** | *Glomeraceae* | *Glomus* | **VTX00334** | *Glomeraceae* | *Glomus* |
| **VTX00137** | *Glomeraceae* | *Glomus* | **VTX00335** | *Paraglomeraceae* | *Paraglomus* |
| **VTX00140** | *Glomeraceae* | *Glomus* | **VTX00337** | *Paraglomeraceae* | *Paraglomus* |
| **VTX00143** | *Glomeraceae* | *Glomus* | **VTX00338** | *Archaeosporaceae* | *Archaeospora* |
| **VTX00149** | *Glomeraceae* | *Glomus* | **VTX00340** | *Claroideoglomeraceae* | *Claroideoglomus* |
| **VTX00151** | *Glomeraceae* | *Glomus* | **VTX00342** | *Glomeraceae* | *Glomus* |
| **VTX00153** | *Glomeraceae* | *Glomus* | **VTX00344** | *Glomeraceae* | *Glomus* |
| **VTX00154** | *Glomeraceae* | *Glomus* | **VTX00347** | *Diversisporaceae* | *Diversispora* |
| **VTX00155** | *Glomeraceae* | *Glomus* | **VTX00348** | *Paraglomeraceae* | *Paraglomus* |
| **VTX00156** | *Glomeraceae* | *Glomus* | **VTX00349** | *Paraglomeraceae* | *Paraglomus* |
| **VTX00159** | *Glomeraceae* | *Glomus* | **VTX00352** | *Paraglomeraceae* | *Paraglomus* |
| **VTX00163** | *Glomeraceae* | *Glomus* | **VTX00354** | *Diversisporaceae* | *Diversispora* |
| **VTX00165** | *Glomeraceae* | *Glomus* | **VTX00356** | *Diversisporaceae* | *Diversispora* |
| **VTX00166** | *Glomeraceae* | *Glomus* | **VTX00357** | *Claroideoglomeraceae* | *Claroideoglomus* |
| **VTX00172** | *Glomeraceae* | *Glomus* | **VTX00358** | *Claroideoglomeraceae* | *Claroideoglomus* |
| **VTX00177** | *Glomeraceae* | *Glomus* | **VTX00359** | *Glomeraceae* | *Glomus* |
| **VTX00186** | *Glomeraceae* | *Glomus* | **VTX00376** | *Archaeosporaceae* | *Archaeospora* |
| **VTX00188** | *Glomeraceae* | *Glomus* | **VTX00380** | *Diversisporaceae* | *Diversispora* |
| **VTX00191** | *Glomeraceae* | *Glomus* | **VTX00399** | *Glomeraceae* | *Glomus* |
| **VTX00193** | *Claroideoglomeraceae* | *Claroideoglomus* | **VTX00401** | *Diversisporaceae* | *Diversispora* |
| **VTX00195** | *Glomeraceae* | *Glomus* | **VTX00402** | *Claroideoglomeraceae* | *Claroideoglomus* |
| **VTX00199** | *Glomeraceae* | *Glomus* | **VTX00409** | *Glomeraceae* | *Glomus* |
| **VTX00202** | *Glomeraceae* | *Glomus* | **VTX00411** | *Glomeraceae* | *Glomus* |
| **VTX00212** | *Glomeraceae* | *Glomus* | **VTX00417** | *Glomeraceae* | *Glomus* |
| **VTX00214** | *Glomeraceae* | *Glomus* | **VTX00419** | *Glomeraceae* | *Glomus* |
| **VTX00219** | *Glomeraceae* | *Glomus* | **VTX00444** | *Paraglomeraceae* | *Paraglomus* |

**Table S2** Overview over VTC (clusters) with number of ASV that were clustered together and possible taxa these VTC belong to, according to taxonomy assigned to the contained ASVs

| **VTC ID** | No of ASVs | SILVA database taxonomy hits (12^th^ rank) | |
| --- | --- | --- | --- |
|  |  | No of taxa | Which taxa (**most assigned sequences**) |
| **VTC00001** | 225 | 11 | *Glomus, Rhizophagus*, **uncultured unclassified**, *Funneliformis,* ***Septoglomus****,* unclassified Glomeromycetes, unclassified Diversisporaceae, unclassified Glomeraceae, unclassified Glomerales, *Claroideoglomus, Otospora* |
| **VTC00002** | 2 | 2 | Unclassified Glomeraceae, *Otospora* |
| **VTC00003** | 2 | 2 | Unclassified Glomeraceae, *Sclerocystis* |
| **VTC00004** | 34 | 1 | *Archaeospora* |
| **VTC00005** | 31 | 1 | *Paraglomus* |
| **VTC00006** | 3 | 1 | *Paraglomus* |
| **VTC00012** | 3 | 1 | *Rhizophagus* |
| **VTC00013** | 1 | 1 | *Septoglomus* |
| **VTC00014** | 4 | 1 | *Septoglomus* |
| **VTC00015** | 2 | 2 | *Archaeospora*, uncultured unclassified |
| **VTC00016** | 1 | 1 | Unclassified Glomeromycetes |
| **VTC00017** | 21 | 4 | Unclassified Glomerales, unclassified Glomeromycota, *Funneliformis,* ***Septoglomus*** |
| **VTC00018** | 15 | 3 | Unclassified Diversisporales, unclassified *Entrophospora*, ***Otospora*** |
| **VTC00019** | 2 | 1 | *Paraglomus* |
| **VTC00020** | 75 | 7 | unclassified Diversisporaceae, ***Diversispora***, unclassified Glomeromycetes, *Funneliformis*, unclassified Diversisporales, *Archaeospora, Claroideoglomus* |
| **VTC00021** | 2 | 1 | *Scutellospora* |
| **VTC00022** | 2 | 1 | Unclassified *Diversispora* |
| **VTC00023** | 3 | 1 | Unclassified *Diversispora* |
| **VTC00024** | 8 | 1 | *Claroideoglomus* |
| **VTC00025** | 7 | 1 | *Claroideoglomus* |
| **VTC00026** | 2 | 1 | *Claroideoglomus* |
| **VTC00027** | 1 | 1 | Uncultured unclassified |

**Table S3** Results of ANOVA of alpha-diversity indices against the history treatment, plant diversity and their interaction. Significant effects (p < 0.05) are shown in bold.

|  | **History treatment** | | **Plant Diversity** | | **Treatment : Diversity** | |
| --- | --- | --- | --- | --- | --- | --- |
|  | **F** | **p** | **F** | **p** | **F** | **p** |
| **VT richness** | 1.00 | 0.371 | 21.02 | **< .0001** | 0.19 | 0.824 |
| ***NH*** | *-* | *-* | 8.25 | **0.005** | - | *-* |
| ***SH*** | *-* | *-* | 6.47 | **0.013** | - | *-* |
| ***PSH*** | *-* | *-* | 10.134 | **0.002** | - | *-* |
| **Shannon** | 5.49 | **0.005** | 7.34 | **0.008** | 0.26 | 0.771 |
| ***NH*** | *-* | *-* | 4.05 | **0.048** | - | *-* |
| ***SH*** | *-* | *-* | 1.24 | 0.269 | - | *-* |
| ***PSH*** | *-* | *-* | 4.53 | **0.037** | - | *-* |
| **Simpson** | 4.78 | **0.009** | 2.64 | 0.109 | 0.138 | 0.872 |
| ***NH*** | *-* | *-* | 1.33 | 0.253 | - | - |
| ***SH*** | *-* | *-* | 0.23 | 0.632 | - | - |
| ***PSH*** | *-* | *-* | 2.20 | 0.142 | - | - |
| **Evenness** | 5.69 | **0.004** | 0.64 | 0.428 | 0.35 | 0.723 |
| ***NH*** | *-* | *-* | 0.65 | 0.422 | - | - |
| ***SH*** | *-* | *-* | 0.02 | 0.889 | - | - |
| ***PSH*** | *-* | *-* | 0.83 | 0.367 | - | - |

**Table S4** Data in φ-coefficients for the two specialised VT and their plant partners.

|  | In treatment | No. of plots that contain | | |
| --- | --- | --- | --- | --- |
|  |  | VT | Plant | Plant and VT |
| VTX00156 (*Glomus*) and  *Gallium mollugo* agg*.* | NH | 25 | 6 | 6 |
| VTX00245 (*Archaespora*) and *Poa pratense* | SH | 16 | 8 | 7 |

**Table S4** Results of ANOVA of plant biomasses, microbial biomass and microbial respiration against a model of history treatment (HT) * plant diversity (Div) and a second more complex model including the plant community composition of functional group diversity (FG), as well as presence/absence of legumes (L), grasses (G) and herbs (H). Significant results (p< 0.05) are bold.

|  | **~ HT * Div** | | | **~ HT + Div + FG + L + G + H** | | |
| --- | --- | --- | --- | --- | --- | --- |
|  |  | **F** | **P** |  | **F** | **P** |
| **Aboveground Plant Biomass** | HT | 10.74 | **< 0.001** | FG | 5.51 | **< 0.01** |
|  | Div | 28.47 | **< 0.001** | L | 4.47 | **0.04** |
|  | HT*Div | 2.66 | 0.07 | G | 0.22 | 0.64 |
|  |  |  |  | H | 3.28 | 0.07 |
| **Legumes** | HT | 0.36 | 0.69 | FG | 1.07 | 0.37 |
|  | Div | 0.69 | 0.41 | L | - | - |
|  | HT*Div | 2.13 | 0.12 | G | 5.81 | **0.02** |
|  |  |  |  | H | 16.87 | **< 0.001** |
| **Grasses** | HT | 3.83 | **0.02** | FG | 2.44 | 0.07 |
|  | Div | 16.90 | **< 0.001** | L | 15.89 | **< 0.001** |
|  | HT*Div | 1.26 | 0.29 | G | - | - |
|  |  |  |  | H | 20.21 | **< 0.001** |
| **Herbs** | HT | 6.06 | **< 0.01** | FG | 0.83 | 0.48 |
|  | Div | 5.11 | **0.03** | L | 0.06 | 0.80 |
|  | HT*Div | 0.81 | 0.44 | G | 26.90 | **< 0.001** |
|  |  |  |  | H | - | - |
| **Root Biomass** | HT | 1.39 | 0.25 | FG | 1.39 | 0.25 |
|  | Div | 15.72 | **< 0.001** | L | 5.37 | **0.02** |
|  | HT*Div | 0.94 | 0.39 | G | 5.03 | **0.03** |
|  |  |  |  | H | 1.15 | 0.29 |
| **Coarse Roots** | HT | 1.39 | 0.25 | FG | 1.77 | 0.16 |
|  | Div | 0.93 | 0.34 | L | 1.59 | 0.21 |
|  | HT*Div | 0.29 | 0.75 | G | 1.59 | 0.21 |
|  |  |  |  | H | 0.32 | 0.58 |
| **Fine Roots** | HT | 0.46 | 0.63 | FG | 1.16 | 0.33 |
|  | Div | 16.37 | **< 0.001** | L | 13.03 | **< 0.001** |
|  | HT*Div | 0.94 | 0.39 | G | 13.09 | **< 0.001** |
|  |  |  |  | H | 1.01 | 0.32 |
| **Microbial Biomass** | HT | 66.26 | **< 0.001** | FG | 2.93 | **0.04** |
|  | Div | 59.45 | **< 0.001** | L | 0.07 | 0.79 |
|  | HT*Div | 10.71 | **< 0.001** | G | 1.11 | 0.29 |
|  |  |  |  | H | 1.55 | 0.22 |
| **Microbial Respiration** | HT | 23.09 | **< 0.001** | FG | 1.49 | 0.23 |
|  | Div | 27.85 | **< 0.001** | L | 1.44 | 0.24 |
|  | HT*Div | 5.53 | **< 0.01** | G | 0.69 | 0.41 |
|  |  |  |  | H | 2.17 | 0.15 |

**Table S5** Results of mediation analyses: Models were run with both the experimental factor history treatment (HT), plant diversity (Div) and the edaphic variants soil P (P) and soil water content (SWC), as well as simpler models with only experimental factors or edaphic variants. Cov1 represents exposure (head of table) and cov2 the outcome (response; left column). Significant results (p < 0.05) are bold. Significance of mediation by AMF equals p-value of cov2.

| **exposure** |  | | **HT + Div + P + SWC** | | **HT + Div** | | **P + SWC** | |
| --- | --- | --- | --- | --- | --- | --- | --- | --- |
| **outcome** |  | **R^2^** | | **p** | **R^2^** | **p** | **R^2^** | **p** |
| **P** | Exp | - | | - | 0.10 | **< 0.001** | - | - |
|  | Out | - | | - | < 0.01 | **0.02** | - | - |
| **SWC** | Exp | - | | - | 0.10 | **< 0.001** | - | - |
|  | Out | - | | - | 0.03 | **< 0.001** | - | - |
| **Plant Biomass** | Exp | 0.14 | | **< 0.001** | 0.10 | **< 0.001** | 0.06 | **< 0.001** |
|  | Out | < 0.01 | | 0.24 | < 0.01 | 0.34 | < 0.01 | **0.04** |
| -        **Legumes** | Exp | 0.16 | | **< 0.001** | 0.12 | **< 0.001** | 0.08 | **< 0.001** |
|  | Out | 0.02 | | **0.04** | 0.02 | **0.04** | 0.02 | 0.03 |
| -        **Grasses** | Exp | 0.15 | | **< 0.001** | 0.12 | **< 0.001** | 0.06 | **< 0.001** |
|  | Out | 0.01 | | 0.26 | 0.01 | 0.12 | < 0.01 | 0.36 |
| -        **Herbs** | Exp | 0.15 | | **< 0.001** | 0.11 | **< 0.001** | 0.07 | **< 0.001** |
|  | Out | < 0.01 | | 0.69 | < 0.01 | 0.64 | < 0.01 | 0.71 |
| **Root Biomass** | Exp | 0.14 | | **< 0.001** | 0.10 | **< 0.001** | 0.06 | **< 0.001** |
|  | Out | < 0.01 | | *0.05* | < 0.01 | *0.05* | 0.01 | **< 0.01** |
| -        **Coarse Roots** | Exp | 0.14 | | **< 0.001** | 0.10 | **< 0.001** | 0.06 | **< 0.001** |
|  | Out | < 0.01 | | 0.73 | < 0.01 | 0.64 | < 0.01 | 0.66 |
| -        **Fine Roots** | Exp | 0.14 | | **< 0.001** | 0.10 | **< 0.001** | 0.06 | **< 0.001** |
|  | Out | 0.01 | | **< 0.01** | < 0.01 | **< 0.01** | 0.02 | **< 0.001** |
| **Microbial biomass** | Exp | 0.14 | | **< 0.001** | 0.10 | **< 0.001** | 0.06 | **< 0.001** |
|  | Out | < 0.01 | | **0.04** | < 0.01 | **0.03** | 0.01 | **< 0.001** |
| **Microbial respiration** | Exp | 0.14 | | **< 0.001** | 0.10 | **< 0.001** | 0.06 | **< 0.001** |
|  | Out | < 0.01 | | 0.11 | 0.02 | **< 0.001** | < 0.01 | **0.02** |
